# Ubiquitin phosphorylation accelerates protein aggregation and promotes neurodegeneration in the aging brain

**DOI:** 10.1101/2023.08.10.552882

**Authors:** Cong Chen, Hua-Wei Yi, Yi Zhang, Tong Wang, Tong-Yao Gao, Zhi-Lin Lou, Tao-Feng Wei, Yun-Bi Lu, Ting-Ting Li, Wei-Ping Zhang, Chun Tang

## Abstract

Ser65-phosphorylated ubiquitin (pUb) was found elevated in neurons of aged and neurodegenerative brains. Yet little is known whether a causative link exists between pUb level and brain aging. Here we show that the knockout of *pink1*, a Ub kinase, abolished pUb elevation and decelerated protein aggregation in aged mouse brains and cells with proteasomal inhibition. Conversely, over-expression of PINK1 but not the kinase-dead version increased the pUb level and accelerated protein aggregation by suppressing of proteasomal degradation. Furthermore, PINK1 over-expression in mouse hippocampus neurons increased pUb level and protein aggregation, slowly leading to mitochondrial injury, neurodegeneration, and cognitive impairment. Notably, the neuronal damages induced by PINK1 were rescued by the dominant negative Ub/S65A mutant, while Ub/S65E phosphomimetic mutant caused neuronal death. Together, an incidental increase of Ub phosphorylation can progressively and cumulatively cause the decline of Ub-dependent proteasomal activity, consequenting promotes neurodegeneration in the aging brain.

## Introduction

Loss of proteostasis is a hallmark of aging and age-related diseases^1^. To maintain a dynamic equilibrium of proteostasis, proteins are constantly synthesized, folded and degraded. Protein degradation involves proteasomal and autophagic-lysosomal processes. Proteasomal degradation occurs to soluble proteins, while the autophagic-lysosomal process usually occurs to large aggregates resistant to proteasomal degradation. Autophagy can also remove damaged organelles, among which mitophagy—the degradation of defective mitochondria— is highly relevant to neurodegeneration and has been well characterized. In neurons, inefficient removal of mitochondria through mitophagy can cause neuronal damage and lead to neurodegeneration^2^.

Proteasomal degradation declines with age throughout the body, which is correlated with increase of protein oxidization, misfolding and ubiquitination^3–5^. Unable to recycle proteins efficiently can lead to the disruption of proteostasis and the formation of protein aggregates. Of note, the non-dividing and long-lived neurons are highly vulnerable to proteostasis disturbance and protein aggregates. Proteasomal degradation is also essential in neuronal activities including synaptogenesis, axon guidance, dendritic spine growth, and learning and memory^6,7^, by processing inhibitory proteins and activating functional proteins involved in PKA^8^, NFκB^9^, CamKII^10^, and other signaling pathways. As such, the decline of proteasomal degradation in neurons contributes to development of neurodegenerative disease and age-related cognitive declines ^6^. Neuronal-specific proteasome augmentation has been shown to extend lifespan and reduce age-related cognitive deficient^11^.

Ubiquitin (Ub) is an important signaling protein in cells, and has been found colocalized with protein aggregates in the brain including Aβ plaques^12,13^. Protein ubiquitination, covalent attachment of Ub or Ub polymeric chain to cellular proteins, is the primary function of Ub, and is installed by activating enzyme E1, conjugating enzyme E2, and Ub ligase E3^14^. When modified with K48-linked poly-Ub, a protein can be targeted to the 26S proteasome for degradation. Proteasomal targeting of the ubiquitinylated protein is achieved through the noncovalent interactions between poly-Ub and one of the Ub receptors, namely Rpn1^15^, Rpn10^16^, and Rpn13^17^, in the proteasome.

Ub itself can be modified^18^. Most notably, Ub is phosphorylated by PINK1 (PTEN-induced kinase 1), a Ser/Thr kinase, at residue Ser65^19,20^. The full-length PINK1 reside at the outer membrane of mitochondria, and the PINK1-Ub-PARKIN axis is the mainly responsible for mitophagy in the event of mitochondrial damages^20–22^. Thus, loss or downregulation of PINK1 activity has been associated with the development of neurodegenerative diseases^19,23^. PINK1 is processed by mitochondrial membrane proteases, affording a 52-kDa shorter version of soluble PINK1 (sPINK1) in the cytoplasm^24,25^. With residue F104 exposed at the protein N-terminus, the sPINK1 is unstable and rapidly degraded by the proteasome in healthy cells^26^. However, the sPINK1 could be stabilized, and the sPINK1 level in the cytoplasm could be elevated in the cultured cells upon pharmacological inhibition of proteasomal activity^27^. Accordingly, the elevation of sPINK1 and pUb has been suggested as an early indicator of the decline of proteasomal degradation^28^.

When phosphorylated, pUb cannot be efficiently utilized by the E1 and E2 enzymes^29^, which would disrupt protein ubiquitination. In neurodegenerative and aged postmortem brains, the pUb level has been shown to increase in neurons^30,31^. The observation led to the proposal that the pUb is a biomarker for damaged mitochondria in aging and mitochondrial damage associated diseases. The increase of pUb in the human autopsy brain has also been associated with α-synuclein pathology in Lewy body disease^32^. Nevertheless, it is unclear whether there is a causative link between the elevation of Ub phosphorylation and the development of neurodegeneration.

Using animal and cell models, here we show that the pUb level increases in naturally aged mouse brains and cells with proteasomal inhibition, whereas inhibiting Ub phosphorylation decelerates protein aggregation. Moreover, we show that an increase of pUb level triggers a progressive decline of Ub-dependent proteasomal degradation, which exacerbate protein aggregation and disrupts proteostasis. In such a vicious cycle that reinforcing one another, the cumulative buildup of pUb would gradually impair mouse neuronal structures and functions, leading to neurodegeneration over an extended time. Thus, elevated pUb level is not only a biomarker and also a risk factor for brain aging and age-related brain diseases.

## Results

### 1. Knockout of *pink1* abolished Ub phosphorylation and alleviated protein aggregation in aged mouse brains

The pUb level increases in the neurons of the aged and diseased human brain^30,31^. We found this is also the case for mouse brain. Using immunofluorescence staining, we found that the pUb level increased in the neurons of the wildtype naturally aged mouse brain (Fig. 1A, top two rows). Using Western blotting analysis, we confirmed that the pUb level is significantly higher by ∼31% in the wildtype aged mouse brain than in the wildtype young mouse brain (Fig. 1B).

**Figure 1.**
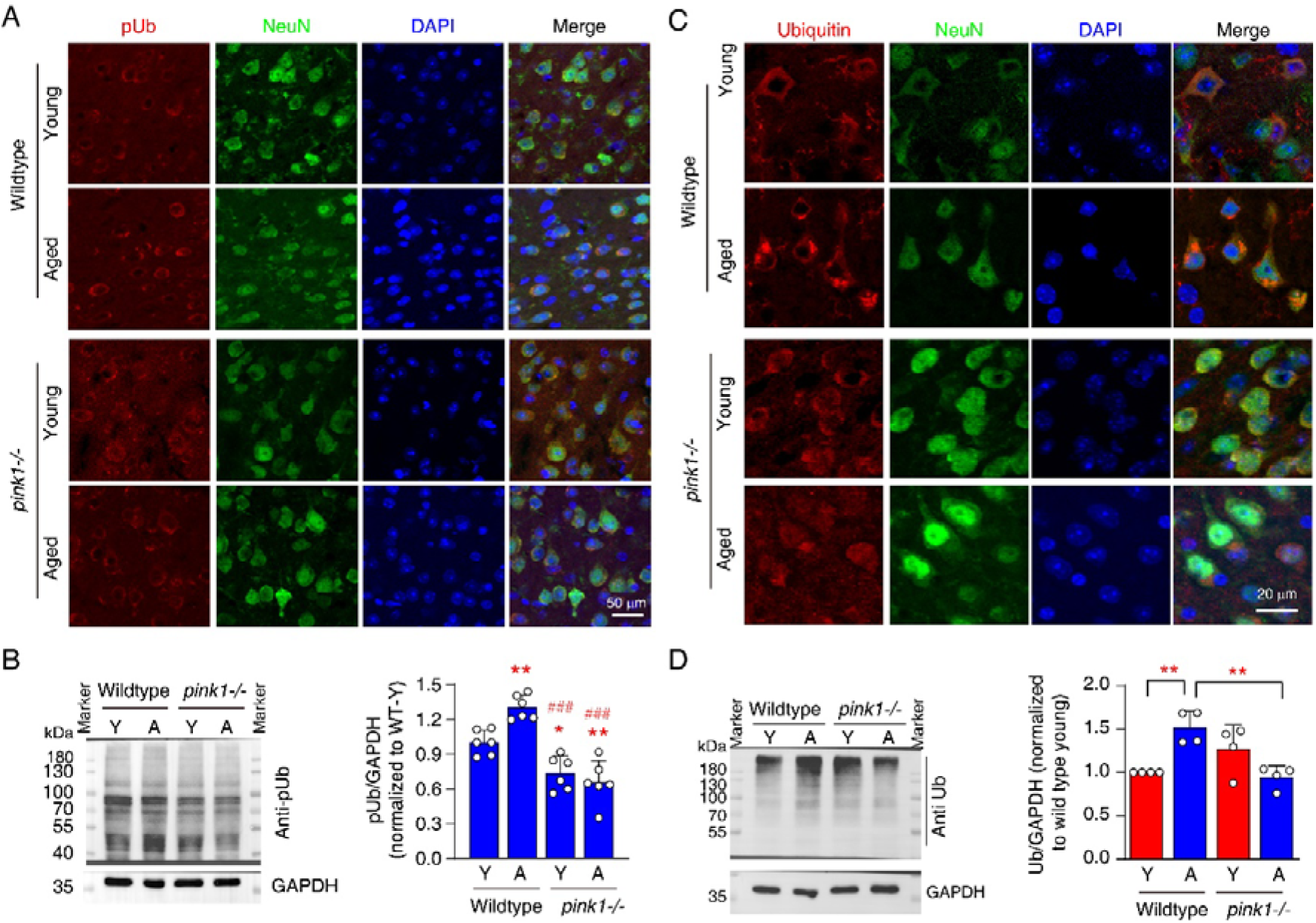
Knockout of *pink1* gene abolished Ub phosphorylation and alleviated protein aggregation buildup in aged mouse brain. **A**. Representative immunofluorescent image of pUb in young (Y) and aged (A) mouse brains. **B**. Western blotting analysis of pUb in young (Y) and aged (A) mouse brain. *N*=6. \**P*<0.05, \*\**P*<0.01, compared to the wildtype young mouse, one-way ANOVA. ^###^*P*<0.001, compared to the wildtype aged mouse, one-way ANOVA. **C**. Representative immunofluorescent image of Ub in young (Y) and aged (A) mouse brains. **D**. Western blotting analysis of Ub in the insoluble fraction obtained from young (Y) and aged (A) mouse brains. *N*=4. \*\**P*<0.01, one-way ANOVA.

PINK1 is the kinase that phosphorylates Ub at residue Ser65. In the *pink1*-knockout (*pink1*-/-) mice, the pUb level remained constant with aging (Fig. 1A, bottom two rows). Western blotting analysis confirmed that there was a significantly lower pUb signal in both young and aged *pink1-/-* mouse brains in comparison to the wildtype (Fig. 1B). The remaining pUb signal in *pink1-/-* mouse sample is likely due to the non-specific binding of anti-pUb antibody, as the band intensities are similar between the young and aged *pink1-/-* mice. Notably, the elevated pUb level in the wildtype aged mouse brain compared to the wildtype young mouse brain should be due to a high PINK1 level. Nevertheless, we could not perform immunofluorescent staining and Western blot analyses for PINK1 due to a lack of working antibodies.

During aging, protein aggregates can build up in neurons due to the decline of proteasomal and autophagy degradation^33,34^. Using an anti-Ub antibody, we observed an even distribution of Ub signal in the cytoplasm of neurons in the wildtype young mouse brain but the formation of puncta in the aged mouse brain (Fig. 1C). Interestingly, in *pink1-/-* mouse, Ub formed puncta in the cytosol of neurons in both young and aged brains (Fig 1C). Using Western blotting analysis, we found that the Ub level significantly increased in the insoluble fraction of wildtype aged mouse brain as compared to that of young mouse brain (Fig. 1D). On the other hand, the Ub level in the insoluble fraction of the *pink1-/-* aged mouse brain is significantly lower than that of the wildtype aged mouse brain (Fig. 1D).

The gradual accumulation of Ub in the insoluble fraction and the Ub-positive puncta in cells are characteristic of protein aggregation. In the aged mouse brain, both pUb and protein aggregation increased, whereas the knockout of *pink1* gene abolished the increase of pUb level and alleviated protein aggregation.

### 2. Knockout of *pink1* abolished Ub phosphorylation and decelerated protein aggregation upon proteasomal inhibition in cultured cells

As sPINK1 is rapidly degraded by proteasomal degradation, we reason that inhibition of proteasomal activity should increase both sPINK1 and pUb levels. We found that the administration of MG132, a potent proteasomal inhibitor, significantly increased the level of sPINK1 but not the full-length PINK1 in HEK293 cells in a concentration- and time-dependent manner, which plateaued after ∼10 hours (Fig. 2A-D). Accordingly, the pUb level also significantly increased in the same fashion as sPINK1 (Fig. 2E-H), but by a smaller extent, manifesting the stochastic nature of protein phosphorylation. As a negative control, the administration of MG132 did not increase either PINK1 or pUb in *pink1-/-* cell (Fig. 2A-H).

**Figure 2.**
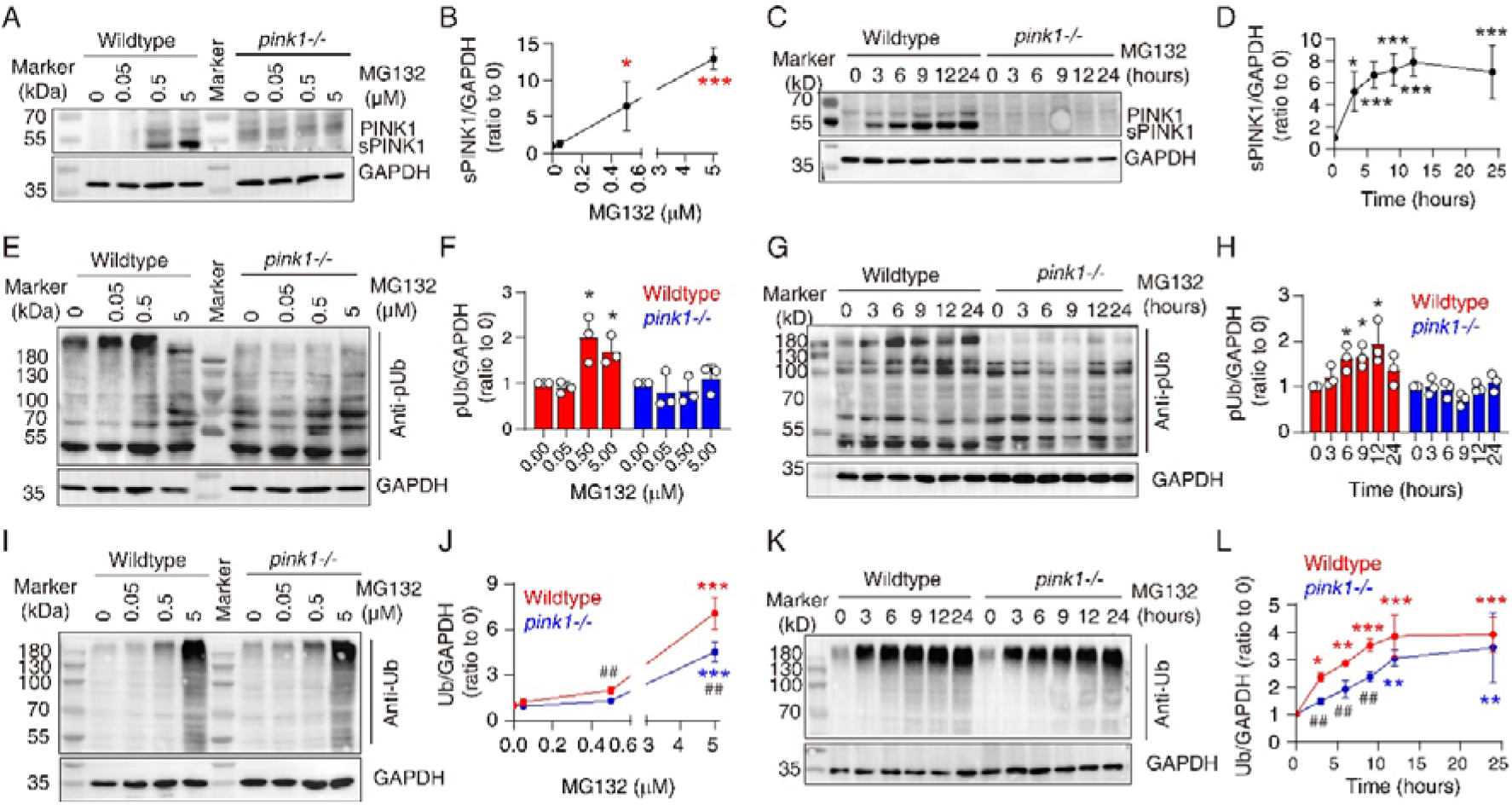
Knockout of *the pink1* gene abolished Ub phosphorylation and alleviated protein aggregation buildup upon proteasomal inhibition in HEK293 cells. **A, B.** Western blotting analysis of PINK1 at 8 hours after administering 0-5 μM MG132 in wildtype and *pink1-/-* cells. *N*=3. \**P*<0.05, \*\*\**P*<0.001, compared with “0” MG132, one-way ANOVA. **C, D.** Western blotting analysis of PINK1 at 0-24 hours after administering 5 μM MG132 in wildtype and *pink1-/-* cells. *N*=4. \**P*<0.05, \*\*\**P*<0.001, compared with time 0, one-way ANOVA. **E, F.** Western blotting analysis of pUb at 8 hours after administering 0-5 μM MG132 in wildtype and *pink1-/-* cells. *N*=3. \**P*<0.05, compared with “0” MG132, one-way ANOVA. **G, H.** Western blotting analysis of pUb at 0-24 hours after administering 5 μM MG132 in wildtype and *pink1-/-* cells. *N*=3. \**P*<0.05, compared with time 0, one-way ANOVA. **I, J**. Western blotting analysis of Ub in the insoluble fraction at 8 hours after administering 0-5 μM MG132 in wildtype and *pink1-/-* cells. *N*=5. \*\*\**P*<0.001, compared with “0” MG132, one-way ANOVA. ^##^*P*<0.01, compared with wildtype, unpaired *t*-test. **K, L**. Western blotting analysis of Ub in the insoluble fraction at 0-24 hours after administering 5 μM MG132 in wildtype and *pink1-/-* cells. *N*=3. \**P*<0.05, \*\**P*<0.01, \*\*\**P*<0.001, compared with time 0, one-way ANOVA. ^##^*P*<0.01, compared with wildtype, unpaired *t*-test.

In parallel to our animal studies, we also evaluated how *pink1* knockout affects protein aggregation in the event of proteasomal inhibition. Using Western blotting analysis, we observed that the treatment of 5 μM MG132 for 8 hours significantly increased the Ub level in the insoluble fraction from HEK293 cell (Fig. 2I, J). In comparison to the wildtype cells, MG132 caused a much smaller increase of Ub in the *pink1-/-* cells (Fig. 2I, J). MG132 at 5 μM significantly increased the Ub level in the insoluble fraction at 3-24 hours after the administration (Fig. 2K, L). In comparison, the accumulation of Ub was significantly lower at 3, 6, and 9 hours upon administering MG132 to *pink1-/-* cells, showing a much slower buildup time course. The Ub level was eventually comparable for the wildtype and *pink1-/-* cells 24 hours after the administration of MG132 (Fig. 2K, L). Thus, the *pink1* gene knockout slowed the protein aggregation, but did not affect the maximal effect of MG132. With no additive effect, the result implies that PINK1 affects the same protein degradation pathway as MG132.

Alternatively, the slowing down of protein aggregate buildup upon the inhibition of proteasome could be due to an enhancement of autophagy. For the wildtype cells, we found the administration of MG132 did increase autophagy in a time- and concentration-dependent manner, as characterized with an increase of LC3 II level (Figure 1SA, B). In comparison, the basal autophagy level is higher in *pink1-/-* cells but decreased upon MG132 treatment (Figure 1SA, B). Thus, we can rule out the enhancement of autophagy as the mechanism that contributes to the alleviation of MG132-triggered protein aggregation in *pink1-/-* cells.

### 3. Over-expression of sPINK1 increased the pUb level and inhibited UPS degradation

We have now shown that proteasomal inhibition caused protein aggregation, accompanied by increased sPINK1 and pUb levels in the cell. Significantly, the knockout of *pink1* gene alleviates protein aggregation. Since Ub is essential in the ubiquitin-proteasomal system (UPS), we then asked whether the elevation of pUb level disturbs UPS pathway for protein degradation.

As the cleavage product of the full-length PINK1, sPINK1 is rapidly degraded per *the N*-end rule, with fully functional proteasomes present in the cell. The pUb level is also in a dynamic flux, as several phosphatases have been identified to remove the phosph^35–37^ oryl group from Ub^35–37^. As a result, the pUb is usually kept at a very low level^38^. To increase the pUb level, we transfected the cells with a mutant sPINK1 (PINK1/F101M/L102-L581, denoted as sPINK1*), which should have a longer half-life than the endogenous sPINK1. We also prepared Ub^GG^-sPINK1 to generate the endogenous sPINK1(F104-L581) in cell^26^. To assess whether the observed effect of sPINK1 is mediated by its kinase activity, we prepared a kinase-dead version of sPINK1* with triple mutations of K219A, D362A, and D384A introduced (sPINK1*-KD)^39^.

After transfecting the full-length PINK1, sPINK1*, and sPINK1*-KD into HEK293 cells, we observed an increase of the respective protein levels using Western blotting analysis (Fig. 3A). Note that a large portion of full-length PINK1 was processed, which gave rise to the 52-kDa sPINK1 band (Fig. 3A). We did not observe the endogenous sPINK1 after the transfection of Ub^GG^-sPINK1 (Fig. 3A), due to its rapid degradation. Accordingly, the pUb level increased after the transfection of PINK1 and sPINK1* (Fig. 3A), but not upon sPINK1*-KD and Ub-GG-sPINK1 transfections (Fig. 3A).

**Figure 3.**
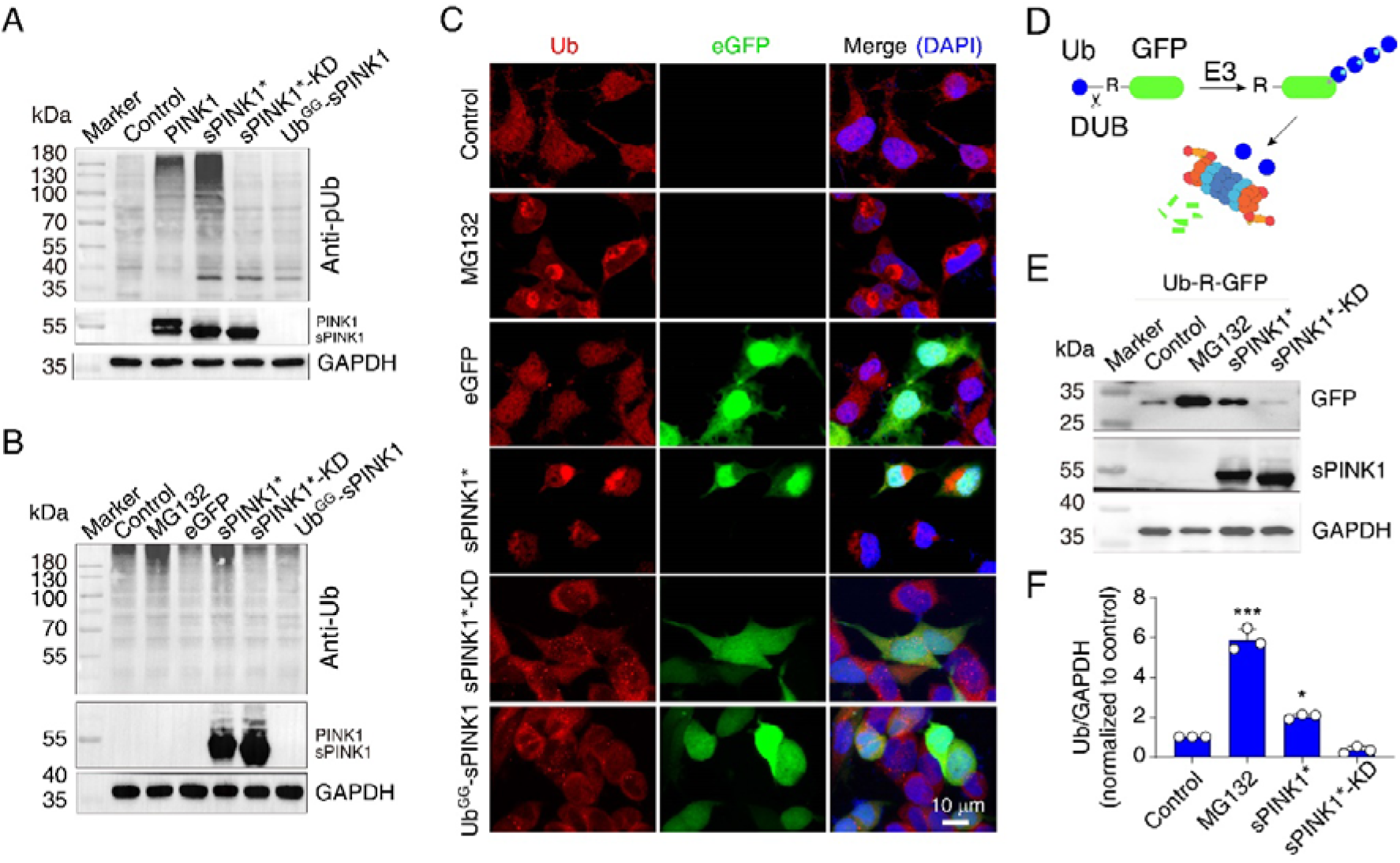
Over-expression of sPINK1 increased pUb level and inhibited Ub-dependent proteasomal degradation. **A**. Western blotting analysis of pUb upon the transfection of various PINK1 constructs. **B**. Western blotting analysis of Ub in the insoluble fractions of cell lysate upon the treatment with 5 μM MG132 or the transfection of various sPINK1 constructs. **C**. Representative immunofluorescent images of Ub staining upon the treatment with MG132 or the transfection of distinct sPINK1. **D**. A schematic for the Ub-R-GFP construct, a model protein for proteasomal degradation. **E.** Representative Western blotting graphs of GFP for assessing Ub-R-GFP degradation in HEK293 cells. **F**. Analysis of the GFP intensity. \**P*<0.05, \*\*\**P*<0.001, compared with control, one-way ANOVA. PINK1 denotes the full-length PINK1. The sPINK1*, the sPINK1* denotes PINK1/Met/F102-L581, the sPINK1*-KD denotes sPINK1*/K219A/D362A/D384A, a kinase dead version, and the Ub^GG^-PINK1(F104-L581) can generate the endogenous sPINK1, F104-L581, after Ub^GG^ is removed in cells.

The treatment of MG132 or the over-expression of sPINK1*, but not the over-expression of sPINK1*-KD and Ub^GG^-sPINK1, increased Ub level in the insoluble fraction obtained from the cultured cells (Fig. 3B). Using immunofluorescent staining, we found that Ub was evenly distributed in the un-transfected and GFP-transfected cells. Importantly, Ub-positive puncta could be observed in the cytoplasm upon the treatment of MG132 or the transfection of sPINK1*, but not upon the transfection of sPINK1*-KD and Ub^GG^-sPINK1 (Fig. 3C). Therefore, protein aggregation is directly related to the elevation of pUb level.

To further uncover the mechanism for the pUb-related protein aggregation, we transfected Ub-R-GFP to the cells and assessed UPS degradation activity (Fig. 3D). The Ub-R-GFP is a model substrate protein of the proteasome, which can be rapidly degraded by fully functional proteasomes^40,41^. Indeed, we found only a low level of GFP could be detected 24 hours post-transfection (1^st^ lane, Fig. 3E, F). MG132 significantly increased the GFP level (2^nd^ lane, Fig. 3E, F). The co-transfection of sPINK1*, but not sPINK1*-KD, rescued Ub-R-GFP from proteasomal degradation (3^rd^ and 4^th^ lane, Fig. 3E, F), though not by as much as MG132 treatment. As such, sPINK1 likely inhibited proteasomal activity by elevating the pUb level. To efficiently modulate the pUb level in the cell, we use sPINK1* but not sPINK1 for the *in vivo* studies.

### 4. Ub phosphorylation inhibited UPS degradation

We then asked how Ub phosphorylation inhibited UPS degradation. The Ub formed di-, tri-, and high-order Ub chains when catalyzed by specific E1 and E2 enzymes (3^rd^ lane, Fig. 4A). In comparison, pUb mainly formed di-Ub when catalyzed by the same enzymes (4^th^ lane, Fig. 4A). The Ub/S65A mutant could be efficiently elongated, affording similar products as non-phosphorylated Ub (5^th^ lane, Fig. 4A); the pUb mimicking Ub/S65E mutant also afford conjugation products of mainly di-Ub (6^th^ lane, Fig. 4A).

**Figure 4.**
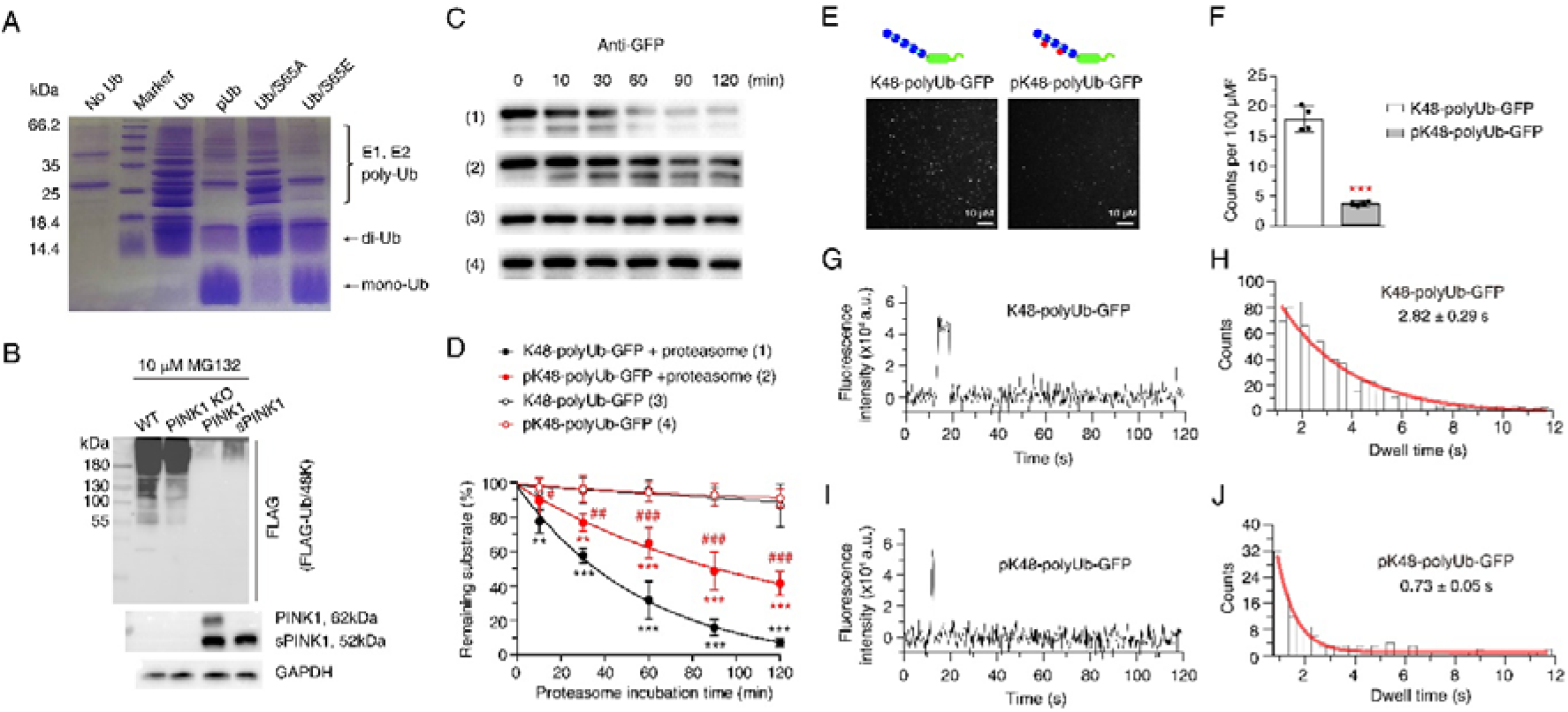
Ub phosphorylation impaired covalent chain elongation and non-covalent interaction with the 26S proteasome. **A**. Assessment of Ub chain catalyzed by E1 and E2 using Coomassie blue staining on the SDS-PAGE. The starting proteins are Ub, pUb, Ub/S65A, and Ub/S65E. **B**. Formation of K48-linked Ub chain in wild-type, *pink1*^-/-^, PINK1-transfected, or sPINK1*-transfected HEK293 cells in the presence of 10 µM MG132. FLAG-Ub/48K was transfected to be elongated and detected. **C**. Western blotting analysis of K48-polyUb-GFP and pK48-polyUb-GFP using an anti-GFP antibody upon *in vitro* proteasomal degradation. **D**. Statistical analysis of the degradation rates of K48-polyUb-GFP and pK48-polyUb-GFP upon *in vitro* proteasomal degradation. *N*=3, ** *p*<0.01, \*\*\**p*<0.001, compared to the respective control without the proteasome degradation; ^#^ *p*<0.05, ^##^ *p*<0.01, ^###^ *p*<0.001, compared to the K48-polyUb-GFP + proteasome. Two-way ANOVA followed with Newman-Keuls multiple comparisons test. **E**. Representative TIRF images of K48-polyUb-GFP (left) and pK48-polyUb-GFP (right) noncovalently interacting with the surface-immobilized proteasome on a glass slide, which would appear as bright puncta. **F**. Analysis of the puncta density. *N*=4, \*\*\**p*<0.001, compared with K48-polyUb-GFP, paired *t*-test. **G, I**. Representative fluorescence traces for a single punctum. **H, J**. Analysis of GFP fluorescence dwell time. The duration of each punctum was binned and fitted using a single exponential decay, shown as the red line.

On the other hand, we assessed the formation of K48-linked Ub chain in HEK293 cells. We transfected the cells with a FLAG-tagged Ub/48K mutant, which could form only the K48-linked Ub chain. We added 10 µM MG132 to preserve the K48-linked Ub chain (1^st^ lane, Fig. 4B). Knockout of *pink1* gene affects little on the length and pattern of the K48-linked Ub chain when determined using an anti-FLAG antibody (2^nd^ lane, Fig. 4B). In comparison, the over-expression of full-length PINK1 and sPINK1* almost obliterated the formation of K48-linked Ub chain (3^rd^ and 4^th^ lanes, Fig.4B). As such, consistent with the previous *in vitro* study^29^, the pUb cannot be efficiently taken up by the E1 and E2 for the post-translational modification of other proteins.

Further, we performed *in vitro* degradation assay using the purified 26S proteasome (Fig. S2). Two ubiquitinated GFP proteins were prepared without or with Ub phosphorylation. We used a recombinant PINK1 to install a phosphoryl group to the Ub chain already conjugated to the GFP, and found that a small fraction of the Ub subunits became phosphorylated (Fig. S3). Upon proteasomal degradation, the ubiquitinated GFP level immediately decreased, and decreased to ∼50% at 30 min (Fig. 4C, D). In comparison, the pUb-modified GFP level only started to drop at 30 min and lowered to 50% at 90 min (Fig. 4C, D).

To explore how Ub phosphorylation inhibited *in vitro* proteasomal degradation, we assessed the interaction between the ubiquitinated substrate protein and the proteasome using total internal reflection fluorescence (TIRF) microscopy. The purified proteasomes were immobilized on a glass slide following the established protocol^42^. Only when the Ub-modified GFP binds to the proteasome for longer than the image-capturing time of the camera, the green puncta from Ub-modified GFP could be visualized. As a negative control, green puncta could not be visualized if the proteasome was not immobilized onto the glass slide or a GFP protein without ubiquitination (Fig. S4). We observed about 5 times the puncta for the proteasome-immobilized slide loaded with K48-polyUb-GFP than with pK48-polyUb-GFP (Fig. 4E, F).

The dwell time of the proteasome-associated substrate protein is inversely related to the protein dissociation rate. The phosphorylated substrate protein dissociated more rapidly from the proteasome than the unphosphorylated K48-polyUb-GFP (Fig. 4G-J, and Fig. S5). Fitting the dwell-time of the visualized GFP puncta, i.e., how long the green puncta were recorded with TIRF microscopy, to a single exponential decay function, we determined the half-life values of the proteasome-substrate complexes at 2.82±0.29 s and 0.73±0.05 s for the K48-polyUb-GFP without (Fig. 4G, H) or with Ub phosphorylation (Fig. 4I, J), respectively. Thus, Ub phosphorylation weakened the interaction between the Ub-attached substrate protein and the proteasome.

Together, we have shown here that the increase of pUb level disrupted the covalent attachment of ubiquitin moieties to a substrate protein and the non-covalent binding to the proteasome, both of which can cause inhibition to UPS degradation activity.

### 5. Proteomics analysis indicated that over-expression of sPINK1* caused neuronal injuries

The pUb level increases in the neurons of the aged brain and the brains with AD and PD^30,31^. Our results here showed that the increase of pUb inhibited UPS degradation. To assess whether the increased pUb level affects neurons, we used an AAV2/9 vector with GFP to specifically over-express sPINK1* in neurons of the hippocampus CA1 region of the mouse brain.

After the transfection of AAV2/9, GFP was co-localized with NeuN immunofluorescence in the hippocampus CA1 region of the mouse brain, indicating successful gene delivery and specific protein expression in neurons (Fig. 5A). Based on the immunofluorescent staining of NeuN, we observed no obvious morphological changes on the 10^th^, 30^th^ and 70^th^ day post-AAV2/9 injection, when compared the groups transfected with either GFP (the control group) or sPINK1* (Fig. 5A). Using Western blotting analysis, we found that sPINK1 significantly increased in the hippocampus of the mouse brain on the 30^th^ day post-transfection (Fig 5B), while the pUb level significantly increased by 45.8% (Fig. 5B, C).

**Figure 5.**
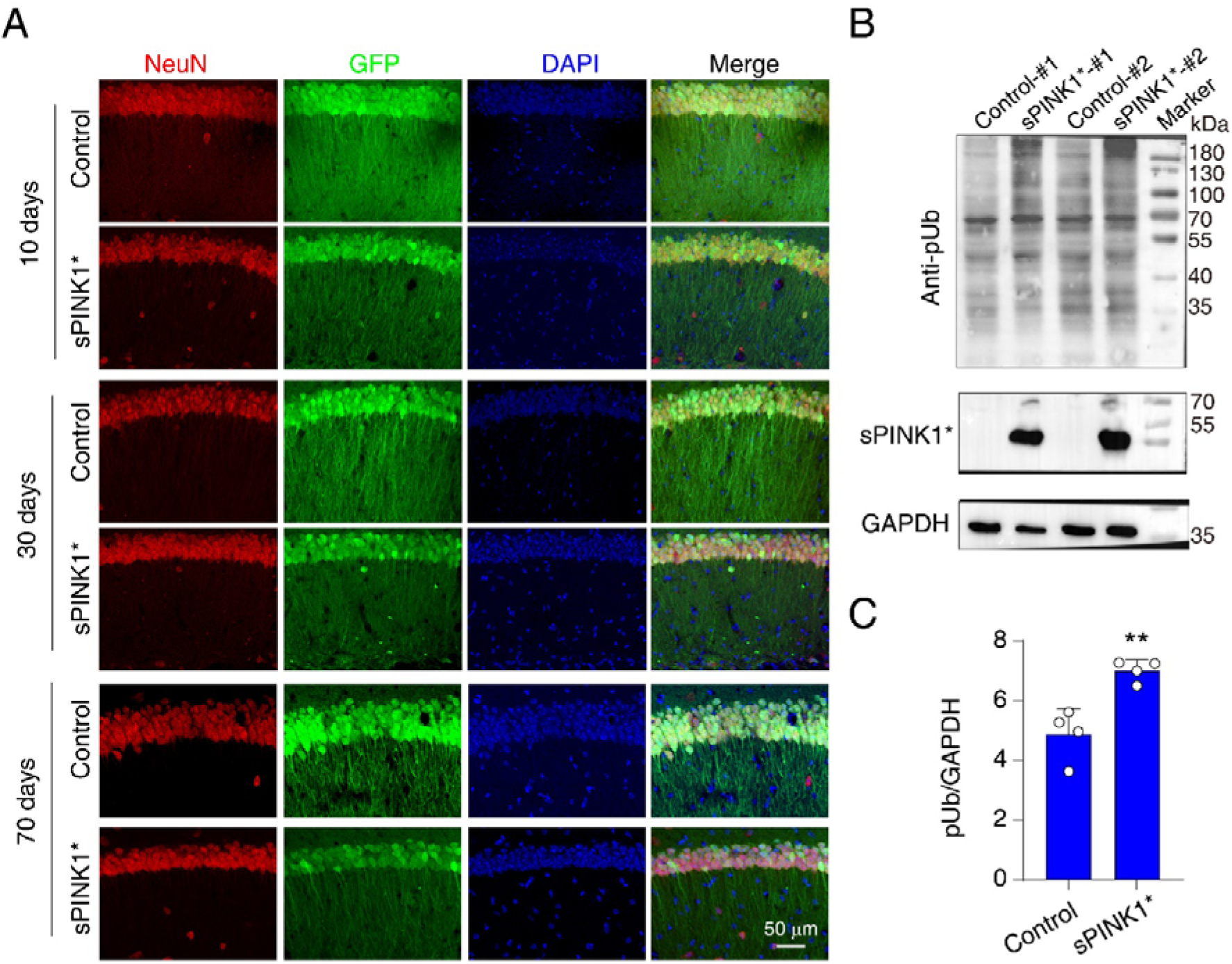
Over-expression of sPINK1* increased pUb level in the hippocampus of the mouse brain. **A**. Representative images of NeuN immunofluorescent staining on the 10^th^, 30^th^, and 70^th^ day post AAV2/9 injection. **B**. Representative Western blotting graphs of anti-pUb and PINK1 on the 30^th^ day post AAV2/9 injection. **C**. Statistical analysis of pUb intensity on the 30^th^ day post AAV2/9 injection. \*\**P*<0.01, compared with control, unpaired *t*-test.

Since the elevation of pUb can inhibit proteasomal degradation that may disrupt proteostasis, we performed proteomics analysis of the hippocampus on the 30^th^ and 70^th^ day after AAV2/9 transfection. The raw data of the change in protein expression level is presented in Table S1. On the 30^th^ day, 4634 proteins were identified in the GFP control group and 4598 proteins in the sPINK1* over-expressing group, with 4128 proteins in common (Fig. 6A). On the 70^th^ day, 4553 proteins were identified in the GFP control group and 4540 proteins in the sPINK1* over-expressing group, with 4108 proteins in common (Fig. 6B). Based on the proteomics data, we performed gene set enrichment analysis (GSEA) for the GO-terms related to mitochondria. On the 30^th^ day post-transfection, significant changes were observed for the proteins related to mitophagy, mitochondrion organization, and respiratory electron transport chain (Fig 6C). On the 70^th^ day post-transfection, however, proteins related to mitophagy became less significant (*p*=0.0804). Significant changes were observed for proteins related to mitochondrial electron transport, tricarboxylic acid cycle, aerobic respiration, apoptotic mitochondrial changes, respiratory electron transport chain, and ATP biosynthetic process (Fig 6C). As such, the over-expression of sPINK1* likely caused mitophagy at the early phase (30^th^ day post-transfection), whereas impairment of mitochondrial function became more significant over time (70^th^ day post-transfection).

**Figure 6.**
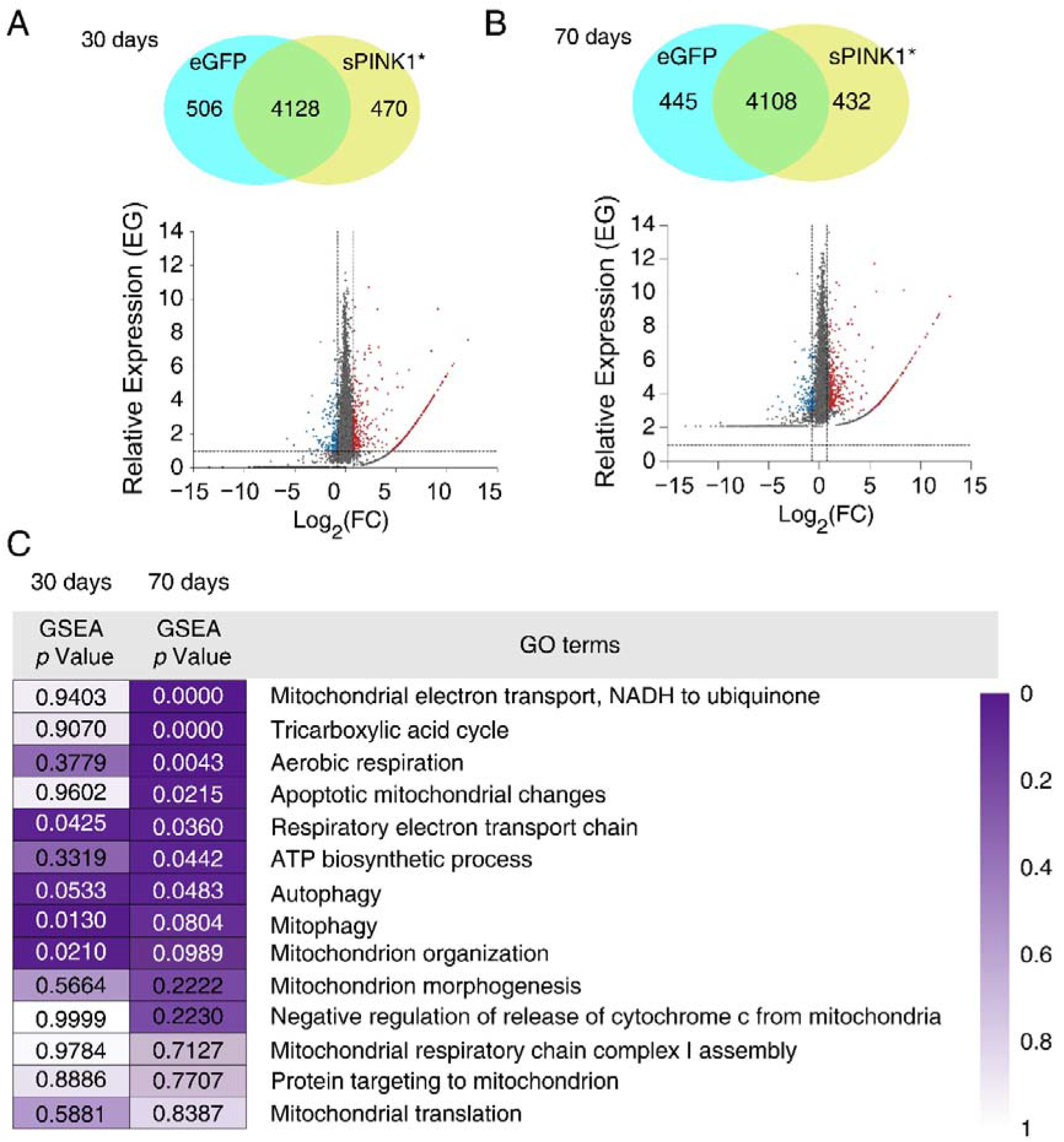
An elevation of the pUb level caused changes in the proteomics of mouse hippocampus on the 30^th^ and 70^th^ day post AAV2/9 injection. **A, B**. The proteomic analysis of the relative expression-FC plot comparing the sPINK1 over-expression and GFP control mouse brain. **C**. The GSEA analysis of mitochondria-related GO terms based on proteomic data.

Based on the proteomics data, we performed gene ontology (GO) analysis and observed widespread proteomic changes (Fig. 7A-F). On the 70^th^ day after AAV2/9 injection into the mouse hippocampus, the over-expression of sPINK1 disrupted proteostasis. Out of the 4985 proteins, we observed 617 proteins were enriched more than two-fold, whereas 624 proteins were halved. The enrichment of the 617 proteins likely resulted from proteasomal inhibition upon sPINK1* over-expression. For example, we observed a significant up-regulation of RNA-binding and RNA-processing proteins, which would otherwise be short-lived^43^. The enrichment can also be a stress response to the insoluble aggregates in cells^44^, as we observed a significant increase of proteins involved in unfolded protein response (Fig. 7A, left). We also observed a significant increase of proteins involved in aging (Fig. 7A, left). The molecular functions of increased proteins were involved in unfolded protein binding, ubiquitin-protein transferase activity, ATPase activity, and calmodulin binding (Fig. 7B, left). Many increased proteins can be assigned to dendrites, dendritic spines, and synapses (Fig. 7C, left).

**Figure 7.**
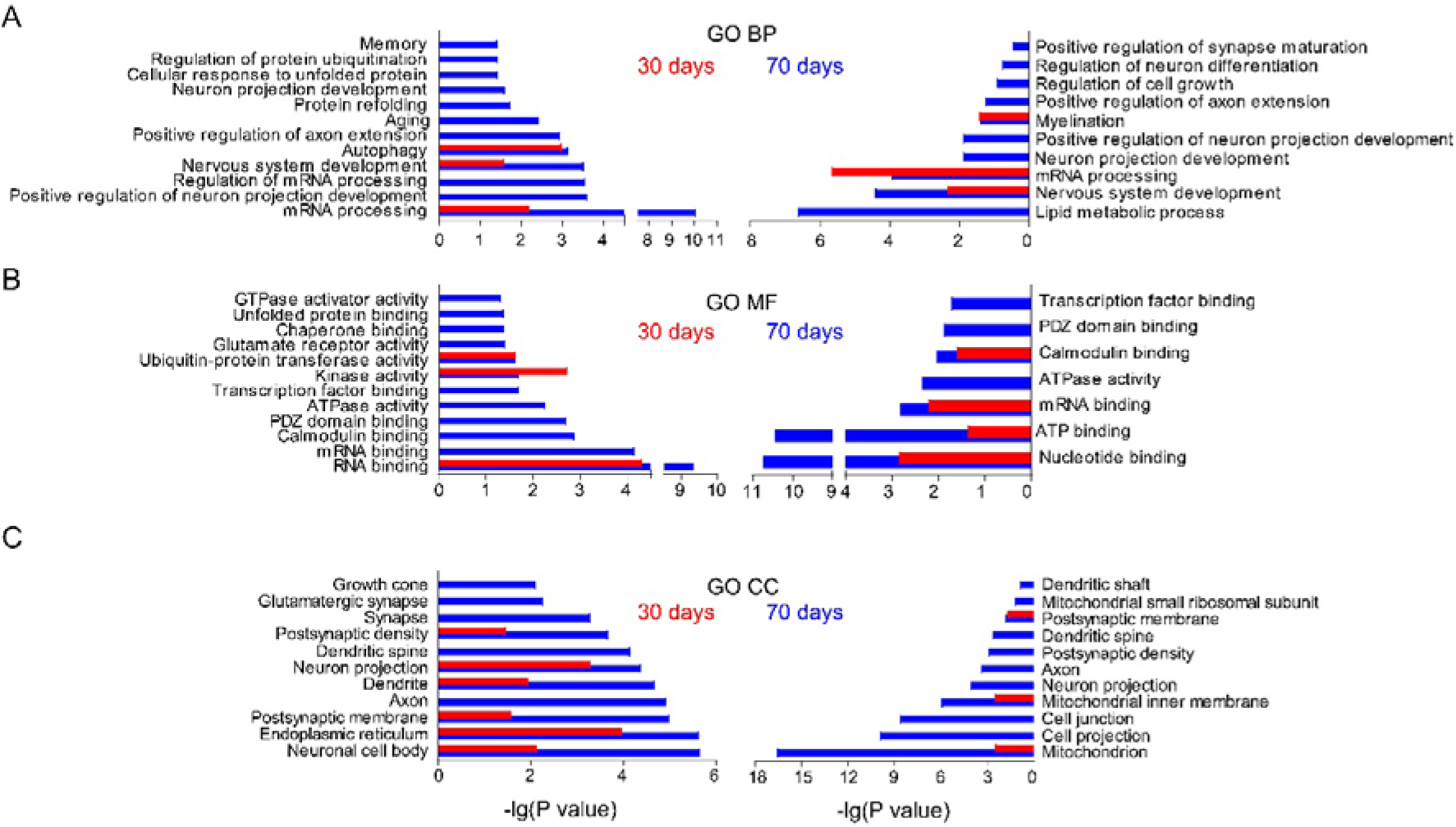
Proteomics analysis revealed an elevation of pUb level altered the GO terms of mouse hippocampus on the 30^th^ and 70^th^ day post AAV2/9 injection. Red bars are analyzed from the proteomics data on the 30^th^ day post AAV2/9 injection, and blue bars, 70^th^ day post AAV2/9 injection. Left panels are analyzed from the 2-fold up-regulated proteins upon sPINK1* over-expression. Right panels are analyzed from the proteins halved upon sPINK1* over-expression. **A**. GO biological process (BP). **B**. GO molecular function (MF). **C**. GO cellular component (CC) analysis of the proteomics data.

On the other hand, the down-regulated proteins are related to myelination, synapse maturation, neuron projection development, and lipid metabolic process (Fig. 7A, right). Functionally, the down-regulated proteins were involved in ATP binding, calmodulin binding, PDZ domain binding, and transcription factor binding (Fig. 7B, right), and can be assigned to the neuronal projection, dendritic spine, and postsynaptic density (Fig. 7C, right image).

As such, the over-expression of sPINK1* not only caused mitochondrial damage, and also exerted a profound and systemic impact on neuronal structure and function. These changes were not as obvious on the 30^th^ day post sPINK1* over-expression as on the 70^th^ day (Fig. 7A-C), suggesting a progressive accumulation of neuronal injuries.

### 6. Elevated pUb level caused protein aggregation in the hippocampus neurons of mice

Based on the proteomics analysis, the over-expression of sPINK1* increased the pUb level on the 30^th^ day post-transfection and injured neurons on the 70^th^ day post-transfection. PINK1 is a kinase with multiple substrates, and the over-expression of sPINK1* could increase the phosphorylation of proteins other than Ub that may also cause neuronal damage. To evaluate the exact role of pUb in this process, we co-expressed Ub/S65A with sPINK1*. The Ub/S65A is a Ub mutant that cannot be phosphorylated by PINK1, and therefore, this dominant negative mutant should block the cellular function mediated by pUb. We also over-expressed a Ub/S65E, which is a mutant that mimics pUb.

First, we characterized the morphological change of neurons upon the co-expression of sPINK1* and Ub/S65A, or the expression of Ub/S65E alone, with the use of immunofluorescent staining of NeuN. We did not observe visible change on the 10^th^, 30^th^, and 70^th^ day following the co-expression of sPINK1* and Ub/S65A (Fig 8A) as compared to the GFP control (Fig 5A). We did not observe morphological changes of the neuron on the 10^th^ day following the expression of Ub/S65E (Fig. 8A). However, the expression of Ub/S65E induced neuronal loss on the 30^th^ and 70^th^ day post the expression (Fig. 8A). Thus, the over-expression of this pUb mimicking mutant is neurotoxic.

**Figure 8.**
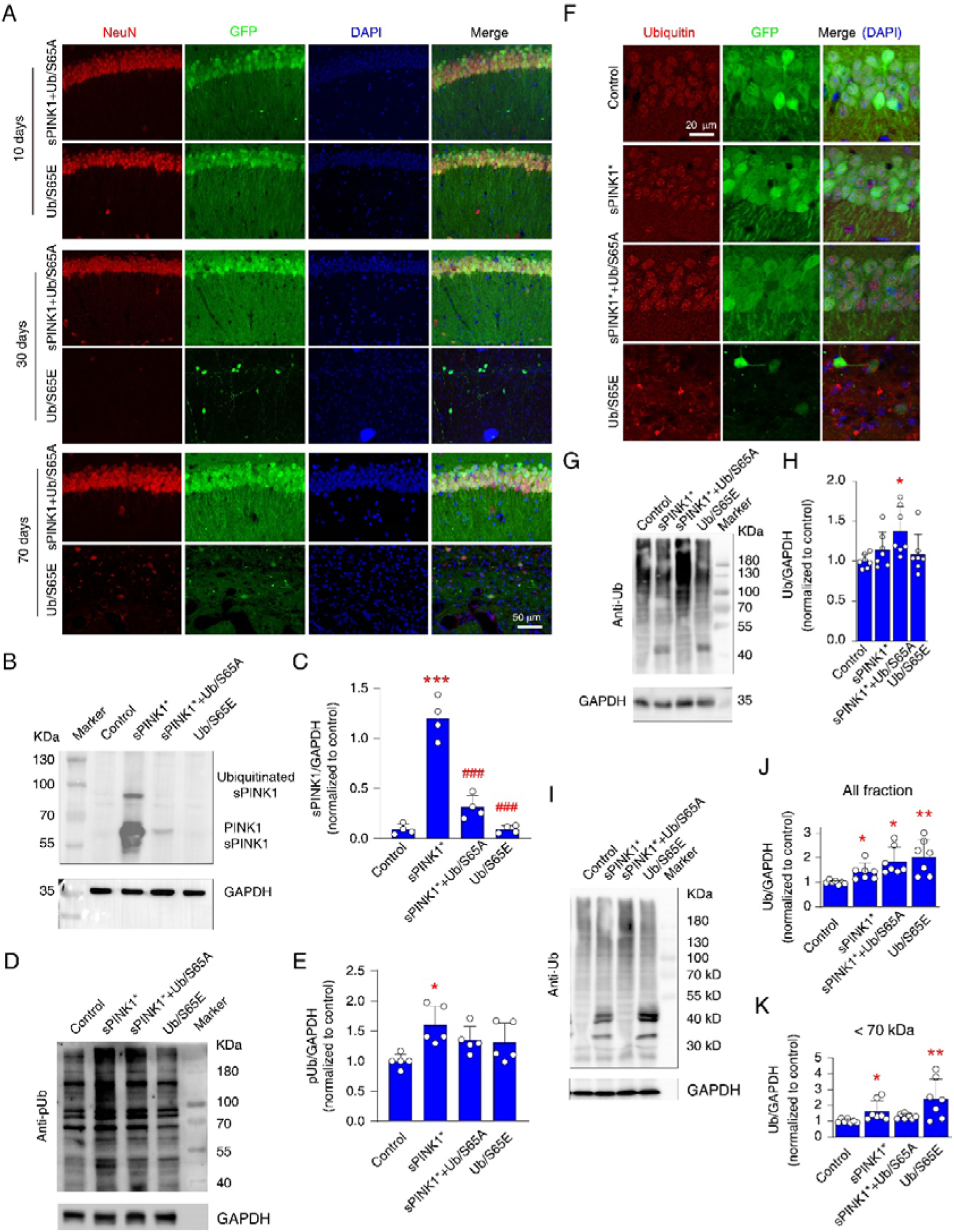
Over-expression of sPINK1* increased pUb level and induced protein aggregation in the mouse hippocampus neuron. **A**. Representative images of immunofluorescent staining of NeuN on the 10^th^, 30^th^, and 70^th^ day post co-expression of sPINK1 and Ub/S65A or over-expression of Ub/S65E. **B, C**. Western blotting analysis of PINK1 on the 70^th^ day post AAV2/9 injection. *N*=4. \*\*\**P*<0.001, compared with control, one-way ANOVA. ^###^*P*<0.001, compared with sPINK1*, one-way ANOVA. **D, E**. Western blotting analysis of pUb on the 70^th^ day post AAV2/9 injection. *N*=5. \**P*<0.05, compared with control, one-way ANOVA. **F.** Representative images of immunofluorescent staining of Ub at the hippocampus CA1 region of the mouse brain. **G, H**. Western blotting analysis of Ub in the soluble fraction of tissue lysis of the hippocampus on the 70^th^ day post AAV2/9 injection. *N*=7, \**P*<0.05, \*\**P*<0.01, compared with control, one-way ANOVA. **I-K**. Western blotting analysis of Ub in the insoluble fraction of tissue lysate of the hippocampus on the 70^th^ day post AAV2/9 injection. The Ub level was quantified in all fractions (J) and the fraction with a molecular weight under 70 kDa (K). *N*=7, \**P*<0.05, \*\**P*<0.01, compared with control, one-way ANOVA.

The transfection of sPINK1* afforded 4 PINK1-derived protein bands on the 70^th^ day (Fig. 8B, lane 2). These bands can be assigned to sPINK1 at 52 kDa, mono-ubiquitinated sPINK1 at around 60 kDa, full-length PINK1 at 63 kDa, and poly-ubiquitinated PINK1 at ∼90 kDa. Quantitative analysis showed that the co-transfection of Ub/S65A dramatically decreased the sPINK1 level (Fig. 8B, C, lane 3), which can be attributed to the rescue of the UPS degradation activity. Accordingly, when compared to the GFP control, the pUb level elevated significantly by 60% upon sPINK1* over-expression (*P*=0.0107) (Fig. 8D). Upon the co-expression of sPINK1* and Ub/S65A, the pUb increased by ∼35% over the GFP control (*P*=0.1927) (Fig. 8E). In fact, considering a much larger pool of both Ub and Ub/S65A in the cell, the effective percentage of pUb over the total Ub can be much lower, which diminished the likelihood of pUb utilization.

Using fluorescent immunostaining of Ub, we observed small puncta in hippocampal neurons of mouse brain upon the over-expression of sPINK1* or the co-expression of sPINK1* and Ub/S65A (Fig. 8F). Large Ub-positive puncta of various sizes were observed upon the over-expression of Ub/S65E (Fig. 8F). Western blotting analysis showed a significant increase of Ub level in the soluble fraction of mouse brain lysate after the co-expression of sPINK1* and Ub/S65A (Fig. 8G, H). In the insoluble fraction of the tissue lysate, the Ub signal was found significantly increased for the sPINK1*, sPINK1*+Ub/S65A, and Ub/S65E groups, as compared to the GFP control, especially for proteins with molecular weights <70 kDa (Fig. 8I-K).

Previously, it has been shown that sPINK1 can phosphorylate p62 and enhance autophagy^27^. Herein we found that the over-expression of sPINK1*, sPINK1*, and Ub/S65A, or Ub/S65E in the mouse brain did not significantly affect the level of LC3-II or p62 in the supernatant of mouse brain lysate (Fig. S6). The level of p62 in the insoluble fraction of protein samples also remained unchanged (Fig. S6). Thus, the autophagic flow is unlikely to be significantly affected by the over-expression of sPINK1* on the 70^th^ post expression, and the increased protein aggregation should mainly result from the decreased UPS activity.

### 7. Elevated pUb level caused mitochondrial damage to hippocampal neurons

We have observed mitochondrial injury on the 70^th^ day post-transfection based on the proteomics analysis (Fig. 6). Using Western blotting analysis, we found that the protein level of PARKIN remained constant upon transfecting different combination of proteins (Fig. 9A, B), yet the pParkin level significantly increased with the transfection of sPINK1* or Ub/S65E (Fig. 9A, C). The co-expression of Ub/S65A largely abrogates PARKIN phosphorylation (Fig. 9A, C). Accordingly, the level of TOM20, a mitochondrial marker, significantly decreased after the over-expression of sPINK1* but was reversed with the co-expression of Ub/S65A (Fig. 10A, D). The over-expression of Ub/S65E also decreased TOM20 level (Fig. 9A, D).

**Figure 9.**
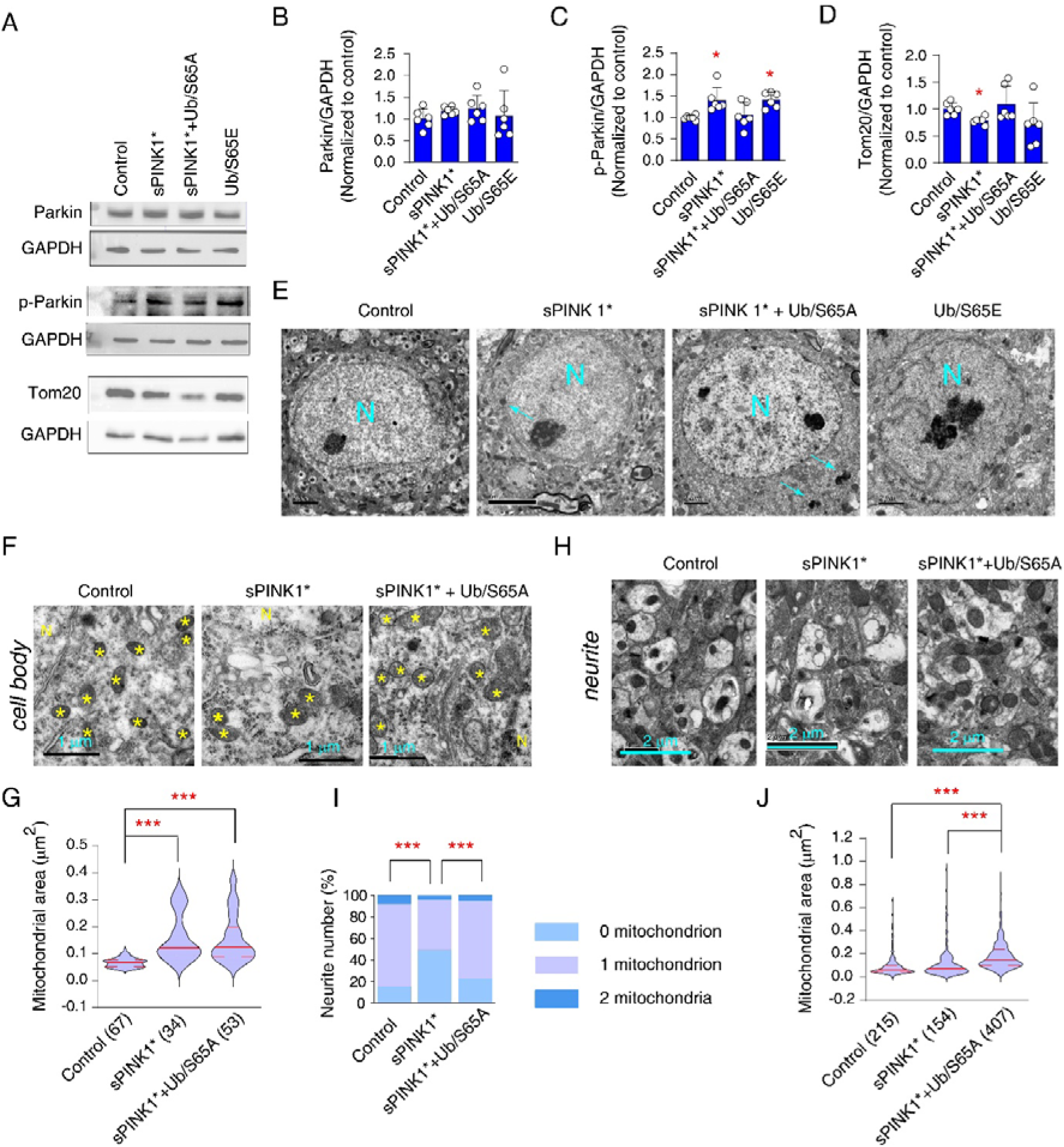
An elevation of pUb level caused the mitochondrial injury of mouse hippocampus neurons. **A-D.** Western blotting analysis of Parkin (A and B), phosphorylated Parkin (pParkin) (A and C), and Tom20 (A and D) in the hippocampus on the 70^th^ day post AAV2/9 injection. *N*=6. \**P*<0.05, compared with control, one-way ANOVA. **E**. Representative transmission electron microscope graphs of hippocampus neurons. “N” represents the nucleus. The cyan arrows indicate lipofuscin plaques. **F**. Representative transmission electron microscope graphs of hippocampus neurons showing the cytoplasmic mitochondria. Yellow stars denote mitochondria. **G**. Statistical analysis of the mitochondrial area within the cytosol of neurons. The number of mitochondria was applied after the group name. The mitochondria were analyzed from 10-13 neurons of 3 mice. \*\*\**P*<0.001, one-way ANOVA. **H**. Representative transmission electron microscope graphs of unmyelinated neurites in the mouse hippocampus. **I.** The neurites are ranked by the total number of mitochondria. \*\*\**P*<0.001, Chi-square test. **J.** Statistical analysis of the mitochondrial area within the neurites. \*\*\**P*<0.001, one-way ANOVA.

**Figure 10.**
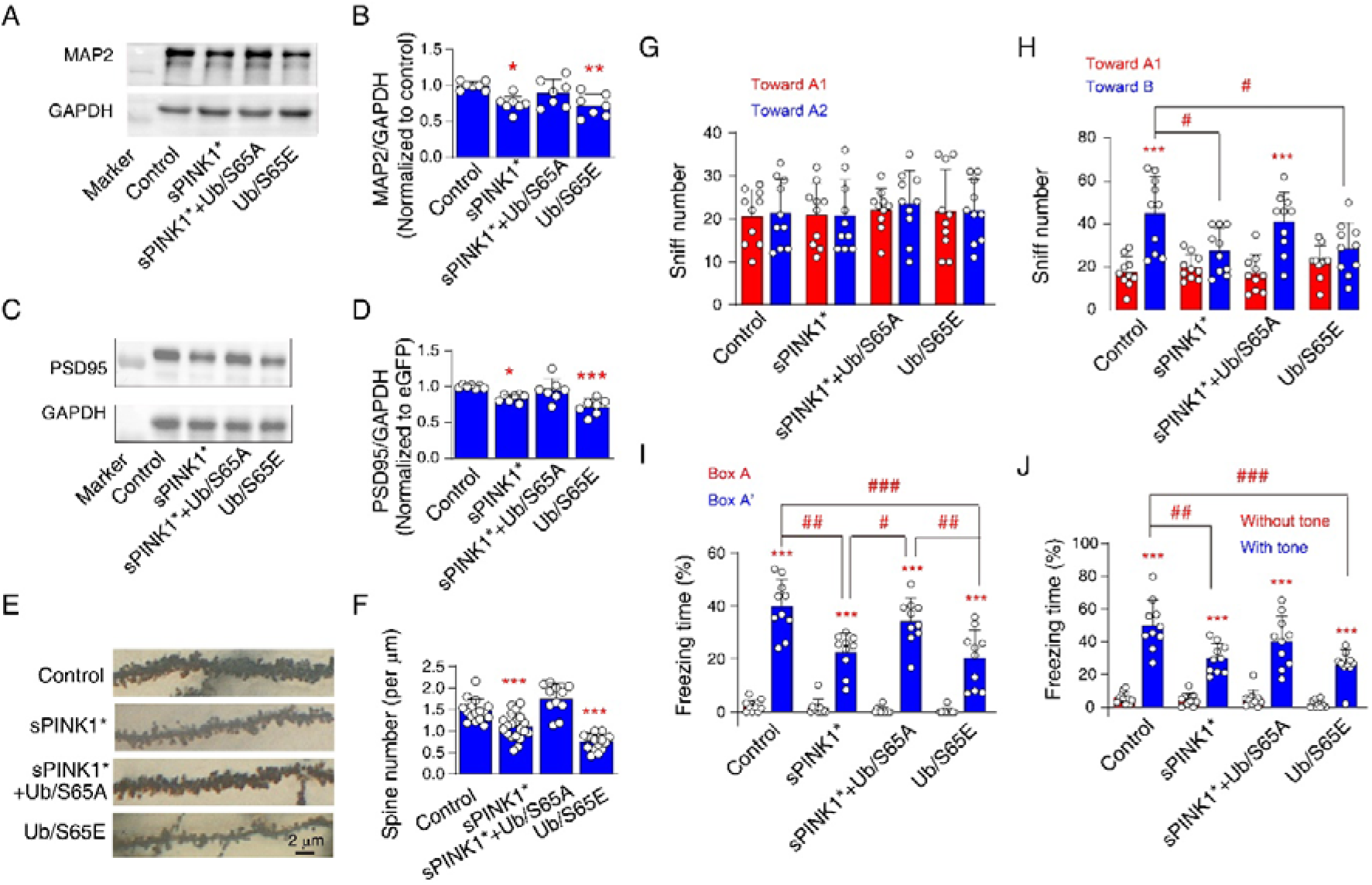
An elevation of the pUb level in the hippocampus injured the neuronal structure and caused cognitive impairment of the mice. **A-D**. Western blotting analysis of MAP2 (A and B) and PSD95 (C and D) in the mouse hippocampus on the 70^th^ post AAV2/9 injection. *N*=7. \**P*<0.05, \*\**P*<0.01, \*\*\**P*<0.001, compared with control, one-way ANOVA. **E.** Representative graphs of Golgi staining from the 2^nd^ branches of hippocampus neurons on the 70^th^ post AAV2/9 injection. **F**. The statistical analysis of the spine intensity in the dendrites of hippocampus neurons. The dendrites were from 3 individual mice. \*\*\**P*<0.001, compared with control, one-way ANOVA. **G.** The sniff number toward two identical objects during the training of mouse novel objective recognition. **H**. The sniff number toward one old object A1 and one novel object A2 during the testing of mouse novel objective recognition. *N*=10. \*\*\**P*<0.001, compared with toward A1 object, paired *t*-test. ^#^*P*<0.05, one-way ANOVA. **I**. The freezing percentage in the same context before and 24 hours after foot shock. *N*=10. \*\*\**P*<0.001, compared with Box A, paired *t*-test. ^#^*P*<0.05, ^##^*P*<0.01, ^###^*P*<0.001, one-way ANOVA. **J**. The freezing percentage in a new box A’ before and after turning on the tone. *N*=10. \*\*\**P*<0.001, compared with without tone, paired *t*-test. ^##^*P*<0.01, ^###^*P*<0.001, one-way ANOVA.

We performed transmission electron microscopy (TEM) to further evaluate mitochondrial injuries. We observed an increased appearance of lipofuscin plaques with high electron density in the mouse hippocampal neurons that over-express sPINK1* or co-express sPINK1* and Ub/S65A (Fig. 9E, S7). Lipofuscin is one of the predominant aggregates in cells and is generally correlated with aging and neurodegeneration^45^. On the other hand, in the Ub/S65E transfected mice hippocampus, we observed apoptotic neurons and necrotic cells (Fig. 9E, S7).

Using the TEM graphs, we analyzed the morphological changes of mitochondria in the cell body and neurites of neurons. The area of mitochondrion within cell body significantly increased after transfecting sPINK1* or sPINK1*+Ub/S65A, compared to the GFP control (Fig. 9F, G). Due to severe neuronal loss, we did not analyze the morphological changes of mitochondria in the Ub/S65E over-expressed mice brain.

The mice with the transfection of GFP had intact neurites, and typically 1-2 mitochondron could be found in the cross-section of the neurites (Fig. 9H, I). The over-expression of sPINK1* decreased mitochondria number in the cross-section (Fig. 9H, I). The co-expression of Ub/S65A reversed the loss of mitochondria in the neurites, with more mitochondria observed in each neurite (Fig. 9H, I). Moreover, the over-expression of sPINK1* increased the mitochondrial area in the neurites from ∼0.064 µm^2^ for the GFP control to ∼0.074 µm^2^. The co-expression of Ub/S65A caused a significant swelling of the mitochondria, affording the medium mitochondrial area of 0.150 µm^2^ (Fig. 9H, J).

Taken together, the elevation of pUb injured mitochondria in neurons and decreased the number of mitochondria by eliciting excessive mitophagy. The co-expression of the Ub/S65A mutant counteracted the effect of sPINK1 and disrupted the mitophagy of the damaged mitochondria, which accounted for the swelling.

### 8. Elevated pUb level altered the structures and functions of hippocampus neurons

The decline of UPS degradation activity and the injuries to the mitochondria can, in turn, profoundly affect the structure and function of neurons. Using Western blotting analysis, we observed that the level of MAP2 (a dendrite marker) and PSD95 (a post-synapse marker) significantly decreased in the mouse brain over-expressing sPINK1* and Ub/S65E, while the co-expression of Ub/S65A restored the protein levels of MAP2 and PSD95 (Fig. 10A-D). We also characterized the dendrite spines using Golgi staining and found that there were fewer spines in the dendrites from the hippocampus neurons of sPINK1* over-expressing mouse brain as compared to the GFP control mouse brain (Fig. 10E, F). The co-expression of Ub/S65A ameliorated the injuries (Fig. 10E, F). On the other hand, the over-expression of Ub/S65E resulted in much slimmer dendrites while significantly decreasing the spine density compared to the GFP control (Fig. 10E, F).

The mitochondrial damage and the change of neuronal structure can further cause cognitive impairment of the mice. Indeed, on the 70^th^ day after transfection, we found that the over-expression of sPINK1* or Ub/S65E impaired novel-object recognition (Fig. 10G, H) and fear conditioning memory (Fig. 10I, J) of the mice. The co-expression of Ub/S65A ameliorated cognitive impairment that was caused by the sPINK1* over-expression (Fig. 10G-J).

## Discussion

In this study, we found that Ub phosphorylation level increases in naturally aged mouse brains, which we show are resulted from the decline of proteasomal activity. The elevation of the pUb level, in turn, causes further inhibition of Ub-dependent proteasomal activity and promotes protein aggregation. For the non-dividing neurons, we found that the elevation of the pUb level led to neurodegeneration, as characterized by mitochondrial injuries, decreased dendritic spines, lipofuscin deposition, and cognitive impairment. After transfecting the Ub kinase sPINK1* into the hippocampus neurons, we found that the neurotoxic effect resulting from the elevation of pUb was much more severe on the 70^th^ day than on the 30^th^ day, indicating a gradual buildup and poor prognosis. Moreover, the neurotoxic effects of pUb correlated with how much pUb or pUb mimic is present. Transfection of Ub/S65E caused massive neuronal death, whereas co-transfection of Ub/S65A alleviated the injuries induced by the sPINK1* over-expression.

PINK1 is a kinase with multiple substrates^46^. The protective effect of Ub/S65A indicates that Ub mediates the neurotoxic effect of sPINK1 instead of other PINK1 substrates. Our results showed that the neurotoxic effect of pUb originated from the inhibition of proteasomal activity, which is distinct from the known effect of pUb on mitophagy. In that well-established role, the pUb exerts a neuroprotective effect by participating in the PINK1-PARKIN axis for the initiation of mitophagy and the removal of damaged mitochondria^20,21,47^.

In the present study, we found that the elevation of pUb did induce mitophagy at an early stage but gradually caused neuronal injuries over time. Consistent with the previous report^29^, pUb is a poor substrate for E1 and E2 enzymes and cannot be efficiently attached to other proteins. Moreover, we show that the pUb chain has a much lower affinity towards the proteasome, shortening the dwell time of ubiquitinated proteins at the proteasome. It has been shown that a sufficiently long dwell time is the prerequisite to commit ubiquitinated proteins for proteasomal degradation^48^. As such, Ub phosphorylation by PINK1 alters covalent Ub polymerization and perturbs the noncovalent interactions between Ub and Ub receptors in the proteasome, both of which would inhibit Ub-dependent proteasomal degradation.

Reciprocally, we show that the elevation of pUb directly resulted from the inhibition of proteasomal activity. The sPINK1 is the cleavage product of full-length PINK1, and is rapidly degraded by the proteasome in healthy cells^26^. Indeed, the pUb level has been found to increase in the aged human brain and in the human AD and PD brains^30,31^, in which the proteasomal activity of neuronal cells is known to be lower^6,49,50^. Thus, a decline of proteasomal degradation leads to a transient spike of pUb, which may further decrease Ub-dependent proteasomal activity. Through this cycle, we show that the pUb is a crucial driver for the progressive decline of proteasomal activity during aging and age-related diseases.

We also show that pUb is neurotoxic at an elevated level. With a progressive decline of proteasomal degradation, the damaged, toxic, and otherwise short-lived proteins accumulate, and insoluble protein aggregates appear in cells. Excessive deposition of protein aggregates can cause mitochondrial damage, leading to neuronal injuries^51–53^. The UPS is also responsible for the removal of endogenous inhibitory proteins of transcription factors in proteins synthesis, including those involved in synapse and neurogenesis ^6^. Pharmacological inhibition of proteasome can slow dendritic spine growth^54^, and impair learning and memory^55^. Based on the proteomics analysis, we found that the increase in the pUb level has a profound effect on the proteins assigned to the dendrite, dendritic spine, and synapse structures. Consequently, the disturbance of proteostasis caused chronic neuronal degeneration and resulted in cognitive impairment of the mice.

The over-expression of Ub/S65E, a positive control of pUb, caused neuronal death, which is much more severe than the over-expression of sPINK1*. One explanation is that the over-expression of Ub/S65E resulted in an equivalent of 100% pUb level, which illustrated a dose-dependent effect of the pUb level. Admittedly, we could not rule out whether other cellular processes also contributed to the observed neuronal injuries. Upon the over-expression of a long-lived version sPINK1, the pUb level increased by only 45.8% and 61% on the 30^th^ and 70^th^ day post-transfection, respectively. Besides the phosphorylation efficiency of the kinase, continuous dephosphorylation of the pUb also accounted for the overall phosphorylation level^35–37^. Considering that the phosphatase activity also declines during aging^56,57^, elevating the phosphatase activity holds the therapeutic promise for exerting neuroprotective effects in the aging brain.

In summary, we have uncovered a novel mechanism for the progressive decline of Ub-dependent proteasomal activity, a hallmark of aging and age-related diseases. The pUb level is elevated upon an inhibition of proteasomal activity, which in turn causes a progressive decline of proteasomal degradation, a gradual buildup of protein aggregates, and eventually neurodegeneration. Thus, increased Ub phosphorylation in the brain is an independent risk factor for the development of aging and neurodegenerative diseases, which warrants pharmacological intervention.

## Methods & Protocols

### Animals

The C57BL/6J male mice in 3, 7-month age were purchased from Zhejiang Academy of Medical Science. The wildtype and *pink1-/-* mice in 18-month were raised in the same cage. Mice were kept in the Laboratory Animal Center, Zhejiang University School of Medicine. The mice had free access to water and food in air-conditioned rooms (20-26 °C, relative humidity around 50%) on a 12-h light/dark cycle. Mice were handled following the Guide for the Care and Use of the Laboratory Animals of the National Institutes of Health. The experimental protocols were approved by the Ethics Committee of Laboratory Animal Care and Welfare, Zhejiang University School of Medicine, with the proven number ZJU20190138.

Using CRISPR-Cas9-mediated genome editing technology^58^, the *pink1* gene knockout mice (C57BL/6J) were customized purchased in Transgenic Mouse Laboratory of Laboratory Animal Center of Zhejiang University. Briefly, two sgRNA sequences that target exon 6 of the *pink1* gene were cloned into pX330-U6-Chimeric_BB-CBh-hSpCas9 plasmid (a gift from Feng Zhang (Addgene plasmid # 42230; http://n2t.net/addgene: 42230; RRID: Addgene 42230) using the *Bbsl* restriction site. The sequence of sgRNA1 is 5’-GACAGCCATCTGCAGAGAGG-3’, and the sequence of sgRNA2 is 5’-GCAGGCAGGACTCACCTCAG-3’. The knockout of *pink1* gene was confirmed by using DAN genotype, quantitative PCR, and Western blotting.

### Injection of recombination Adeno-Associated Virus (rAAV) to mouse hippocampus CA1

The rAAV-EF1a-WPRE-hGH pA (AAV2/9) was used to specifically express proteins within neurons. The rAAV-EF1a-EGFP-WPRE-hGH-pA can express GFP and is a virus infection control. The rAAV-EF1a-sPINK1-P2A-EGFP-WPRE-hGH pA can express GFP and sPINK1 (PINK1/(M102-581) as two separate proteins. The rAAV-EF1a-Ub/S65A-P2A-sPINK1-P2A-EGFP-WPRE-hGH pA can express GFP, sPINK1, and Ub/S65A as three separate proteins. The rAAV-EF1a-Ub/S65E-P2A-EGFP-WPRE-hGH pA can express GFP and Ub/S65E as separate proteins.

Mice with 7-months old were fixed on a mouse stereotaxic holder (RWD Life Science Co. LTP, China) for the injection of AAV2/9. The distinct AAV2/9 was stereo-tactically injected bilaterally into the CA1 region using a microinjector (Hamilton syringe) with a 31-gauge needle and micropump (LEGATO® 130 SYRINGE PUMP, KD Scientific Inc). After anesthetized using pentobarbital sodium (100 mg/kg, ip), the injections were made at stereotaxic coordinates of Bregma: anterior-posterior (AP) = −1.7 mm, mediolateral (ML) =1.5 mm, dorsoventral (DV) = 1.7 mm. A total 0.3μl AAV2/9 (2.25E+12 viral particles/ml) was injected into each side of the hippocampus.

### Behavioral tests

Novel object recognition (NOR) was used to evaluate the spatial memory of mice according to our previous report^59^. After handling for 5 days, the test was performed on the 68^th^ day after the intra-hippocampus injection of the virus. An open-field test system (ViewPoint Behavior Technology, France) was used for NOR detection. During training, two identical objects (named object A1 and A2, 5 cm × 5 cm × 5 cm blue cone) was placed diagonally. The mouse was put at the center of the box, and the movement of the mouse was recorded for 10 min. After returning to the home cage for 24 hours, the mouse was put in the same box with one of the objects replaced by an object in distinct color and shape (object B, 5 cm × 5 cm × 10 cm yellow cuboid). The replaced object was assigned as a novel object. The sniff number (number of visits) toward the objects was analyzed.

Fear conditioning test was performed in the apparatus (ACT-100A, Coulbourn Instruments Inc., Lehigh Valley, PA, USA). The mouse was put into a context (named Box-A) for 2 min, then an 85 dB, 3 kHz tone was activated for 30 sec. Two seconds before the end of the tone, a 2 seconds foot shock was applied (1 mA). After foot shock, the mice were kept in BoxA for another 30 sec, and they were returned to the home cage. Twenty-four hours after training, the mice were put back to the Box-A, and allowed freely explore for 5 min. The freezing time in Box A was measured, which indicates contextual memory. After the mice were put in the home cage for 2 hours, they were placed in a new context (named Box-B), and allowed freely explore for 2 min. Then the tone accompanied with foot shock was applied for 3 min. The freezing time in Box-B with the tone indicates cue-memory.

### Mouse tissue collection

Ten, 30, or 70 days after the AAV2/9 injection, mice were anesthetized using pentobarbital sodium (150 mg/kg, ip). The mouse brains were removed after transcardially perfused with 4 °C saline and freshly prepared 4% paraformaldehyde. The brains were further fixed in 4% paraformaldehyde at 4 °C for one day and transferred to 30% sucrose for three days for dehydration. The brains were sliced into 30 μm thick slices using cryo-microtomy (CM1900, Leica, Wetzlar, Germany). The slices were stored at −20°C in the solution (30% glycerol, 30% glycol, 40% PBS) for immunofluorescent staining.

For the Western blotting analysis, the mouse was transcardially perfused only with 4 °C saline. The brains were removed, and the hippocampus was isolated. The hippocampus was quickly frozen in liquid nitrogen and stored at −70 °C until use.

### Cell lines, cell culture and plasmid transfection

HEK293 cells were purchased from the Institute of Cell Biology of the Chinese Academy of Sciences (Cell Biology of the Chinese Academy of Sciences, Shanghai, China). The *pink1* gene knockout HEK293 cell was homely made using CRISPR-Cas9-mediated genome editing technology^60^. PINK1 sgRNA (CCTCATCGAGGAAAAACAGG) was cloned to U6-sgPINK1-mCherry plasmid (a gift from John Doench & David Root (Addgene plasmid # 78038; http://n2t.net/addgene:78038; RRID:Addgene_78038)). The U6-sgPINK1-mCherry plasmid and PX458-Cas9-EGFP plasmid (a gift from Prof. Feng Zhang (Broad Institute, Cambridge, MA), Addgene plasmid #48138) were co-transfected into HEK293 cells. The mCherry and GFP double-positive cells were single-cell sorted 24 h post-transfection using an MoFloAstrios EQ cell sorter (Beckman Coulter, US) and grown in separate cultures that were subsequently screened for the presence of frameshift mutations leading to nonsense-mediated decay on both alleles. PINK1 KO was confirmed using western blotting.

Cells were grown in Dulbecco’s modified essential medium (DMEM, Gibco by Thermo Fisher Scientific, C11995500BT) supplemented with 10% heat-inactivated fetal bovine serum (FBS, Zhejiang Tianhang Biotechnology, China, 11011-8611) in the atmosphere of 5% CO_2_ under 37 °C.

Lipo3000 (Invitrogen, California, USA, L3000001) was used to transfect plasmids, according to the manuscript. Briefly, a suitable number of cells were seeded one day before transfection with around ∼70-80% confluent at the time of transfection. Before transfection, plasmid DNA was carefully mixed with 250 μl Opti-MEM (Gibco by Life Technologies, Carlsbad, USA). Then 250 μl Lipo3000 was carefully mixed with Opti-MEM. The mixture was incubated at room temperature for 25 min. The freshly formed DNA/Lipo3000 precipitates were carefully pipetted to the cells. Six hours later, the medium containing transfection reagents was removed, and fresh medium was added.

For pharmacological inhibition of proteasome, cells were treated with 0, 0.05, 0.5, 5 μM MG132 for 8h, or treated with 5 μM MG132 for 0, 3, 6, 9, 12, 24 hours.

At the end of treatments, the cells on cover slides were washed with 37°C PBS and then fixed using freshly prepared 4% paraformaldehyde. Then, they were stored in PBS under 4 °C for immunofluorescent staining. For western blotting, cells were collected using trypsin digestion after washing twice using PBS.

### The construction of Ub-R-GFP vector

The Ub-R-GFP is used for the *in vivo* measurement of proteasomal degradation activity, and constructed according to a previous report^40^. Briefly, the ubiquitin open reading frame was amplified by PCR with the following primers:

Sense primer: 5’-GCG GAATTCACCATGCAGATCTTCGTGAAGACT-3’

Antisense primer: 5’-GCG GGATCCTGTCGACCAAGCTTCCCGCGCCCACCTCTGAGACGGAGTAC-3’

The PCR product was cloned into the EcoRI and BamHI sites of the EGFP-N2 vector. Beside the R residue, a 12 amino acid peptide was inserted between Ub and GFP to increase the proteasomal degradation. Thus, the final product is Ub-R-GKLGRQDPPVAT-GFP.

### Immunofluorescence staining

The floating brain slices or cells on cover glasses were incubated in 0.1% Triton-X PBS for 30 min, and followed by the incubation of 5% donkey serum for 1hr. Then the samples were incubated with mouse anti-ubiquitin antibody (1:200, Santa Cruz Biotechnology, Texas, USA; SC-8017), rabbit anti-pUb antibody (1:200, Millipore, Massachusetts, USA; ABS1513), rabbit anti-PINK1 antibody (1:200, Novus, Colorado, USA; BC100-494), rabbit anti-NeuN antibody (1:1000, CST, Pennsylvania, USA; #36662) at 4 °C overnight. After washed with PBS (10 min for 3 times), the samples were incubated with Cy™3 AffiniPure Donkey Anti-Mouse IgG (H+L) (1:200, Jackson ImmunoResearch, PA, USA;715-605-150) or Cy™3 AffiniPure Goat Anti-Rabbit IgG (H+L) (1:200, Jackson ImmunoResearch, PA, USA;715-605-150; 111-165-003) or Alexa Fluor® 488 AffiniPure Goat Anti-Rabbit IgG (H+L) (1:200, Jackson ImmunoResearch, PA, USA; 111-545-003) or Alexa Fluor® 488 AffiniPure Goat Anti-Rabbit IgG (H+L) (1:200, Jackson Immuno Research, PA, USA; 715-545-150) for 2 h. After washing with PBS (10 min’ 3 times), the slices were mounted on slides using a ProLong™ Gold Antifade Mountant with DAPI (Invitrogen Corp., Carlsbad, CA, USA). The images were taken under an Olympus FV100 confocal microscope (Olympus, Japan) and analyzed using MetaMorph Offline (version 7.8.0.0, Molecular Devices, LLC. San Jose, CA 95134 USA

### Western blotting

The cells for the Western blot detection were collected using trypsin after washing twice using PBS. Then RIPA lysis buffer (Beyotime Biotechnology Research Institute, Jiangsu, China; P0013B) with protease inhibitor (Beyotime Biotechnology Research Institute, Jiangsu, China; P1005) and phosphatase inhibitor (Beyotime Biotechnology Research Institute, Jiangsu, China; P1081) was applied. The cells were cracked on ice for 30 min. Every 10 min, the cells were vortexed. Then, the cell samples were centrifuged at 12000 rpm for 30 min under 4 °C. The supernatant was collected as the soluble protein sample for Western blotting analysis. The precipitation was resuspended in 20 μl SDS buffer (2% SDS, 50 mM Tris-HCl, pH7.5) for ultrasonic pyrolysis under 4 °C. The ultrasonic pyrolysis cycle included 10 sec for ultrasonic pyrolysis (Diagenode, Seraing, Belgium) and 30-sec intervals. And a total of 8 cycles was applied until there was no precipitation. Then the samples were centrifuged at 12000 rpm for 30 min under 4 °C. The supernatant was collected as the insoluble protein sample for Western blotting analysis.

For the sample preparation from mouse hippocampus, the mice were anesthetized by intraperitoneal injection of pentobarbital sodium (150 mg/kg) before sacrifice. The brains were quickly removed after transcardially perfused with 4 °C saline. After adding RIPA Lysis Buffer (0.5 ml/100mg tissues) with protease inhibitor and phosphatase inhibitor, the brain samples were thoroughly homogenized with a precooled Tissue Prep instrument (TP-24, Gering Instrument Company, Tianjin, China) for 1 min at 4 °C. After centrifuging at 12000 rpm for 30 min under 4 °C. The supernatant was collected as the soluble protein sample. The precipitation was resuspended in 20 μl SDS buffer (2% SDS, 50 mM Tris-HCl, pH7.5) for ultrasonic pyrolysis under 4 °C. The ultrasonic pyrolysis cycle included 10 sec for ultrasonic pyrolysis (Diagenode, Seraing, Belgium) and 30-sec intervals, a total of 8 cycles were applied until there was no precipitation. Then the samples were centrifuged at 12000 rpm for 30 min under 4 °C. The supernatant was collected as the insoluble protein sample for Western blotting analysis.

All protein concentration was determined by using BCA Protein Assay Kit (Beyotime Biotechnology, Shanghai, China; P0009). Appropriate protein samples (50-100 μg) were used for Western blotting analysis. The following antibodies were used: mouse anti-ubiquitin antibody (1:800, Santa Cruz Biotechnology, Texas, USA; SC-8017), rabbit anti-pUb antibody (1:1000, Millipore, Massachusetts, USA; ABS1513), rabbit anti-PINK1 antibody (1:1000, Novus, Colorado, USA; BC100-494), mouse anti-GAPDH antibody (1:5000, Proteintech, Wuhan, China; 60004-1-Ig), mouse anti-MAP2 antibody (1:2000, Millipore, Billerica, MA, USA; AB5622), rabbit anti-PSD95 antibody (1:1000, CST, Pennsylvania, USA; 3450S), mouse anti-Parkin antibody (1:1000, CST, Pennsylvania, USA; 4211S), rabbit anti-pParkin antibody (1:1000, CST, Pennsylvania, USA; 36866S), rabbit anti-Tom20 antibody (1:1000, Pennsylvania, USA; 42406S), rabbit anti-LC3 antibody (1:1000, Sigma, Massachusetts, USA; L7543), rabbit anti-p62 antibody (1:1000, Abcam, California, USA; ab109012), mouse anti-FLAG antibody (1:10000, TransGen biotech, Beijing, China; HT201-01). The secondary antibody was HRP-conjugated goat anti-mouse IgG (1:3000, Cell Signaling Technology, MA, USA; 7076S) or HRP-conjugated goat anti-rabbit IgG (1:10000, Jackson Immuno Research, PA, USA, 111-035-003). The immunoblots were then detected using ECL reagents (Potent ECL kit, Multi Sciences Biotech, Hangzhou, China; P1425) and measured by using GBOX (LI-COR, Odyssey-SA-GBOX, NE, USA). The results were normalized to GAPDH (as a loading reference) or Ponceau staining (Beyotime Biotechnology Research Institute, Jiangsu, China; P0022), and then normalized to the eGFP-expression control on the same immunoblot membrane.

### Transmission electron microscopy

Seventy days after AAV2/9 administration, mice were anesthetized by intraperitoneal injection of pentobarbital sodium (150 mg/kg) before sacrifice. The mice were transcardially perfused with 4 °C saline, and freshly prepared 4% paraformaldehyde. The dorsal hippocampus was separated and fixed in 2.5% glutaraldehyde for 12 hours at 4 °C.

The samples were washed three times using PBS and fixed in 1% osmic acid at 4 °C for 1 hour. After rinsing with water three times, the samples were pre-stained in 4% uranyl acetate at 4 °C for 30 min. Then the samples were sequentially dehydrated in 50%, 70%, 90%, 100% alcohol, and 100% acetone at 4 °C for 20 min. The samples were then infiltrated in 1:1 acetone-EPON812 (Araldite, Electron Microscopy Sciences, Hatfield, PA, USA) for 2 h at room temperature and in 100% EPON812 for 2 h at room temperature. Samples were polymerized at 37 °C for 24 h, 45 °C for 24 h, and 60 °C for 48 h. After polymerization, 120 nm thin sections were cut on an Ultracut microtome (Leica, Germany). The sections were stained in 4% uranyl acetate for 20 min and 5 min in lead citrate, rinsed, and dried.

Images were taken using an 80 keV Philips TECNAI 10 transmission electron microscope equipped with Ganta 794 CCD.

### Golgi staining

Seventy days after AAV2/9 administration, mice were anesthetized by intraperitoneal injection of pentobarbital sodium (150 mg/kg) before sacrifice. The mouse brain was quickly removed, and the dorsal hippocampus was separated. The hippocampus was immediately fixed in the fixative (Servicebio, Wuhan, China, G1101) for more than 48 hrs.

Cut the dorsal hippocampus with a thickness of 2-3 mm thickness around the injection site. Gently rinse the tissue with normal saline several times. Place the hippocampus tissue into Golgi-cox staining solution (Servicebio, Wuhan, China; G1069), and incubate in a cool and dark place for 14 days. The Golgi-cox staining solution was changed 48 hours after the soak, and change the new staining solution every 3 days. Fourteen days after staining, the tissues were immersed in distilled water 3 times and incubated in 80% glacial acetic acid overnight until the tissue became soft. After rinsing with distilled water, the tissue was placed into 30% sucrose.

Cut the tissue into 100 μm with an oscillating microtome, and place the slices on a gelatin slide, dry overnight in the dark. Then treat the slide with ammonia water for 15 min. After washing with distilled water, the slide was incubated in the fixing solution for 15 min. Finally, after washing, the slide was sealed using glycerin gelatin, and images were taken using VS120 Virtual Slide Microscope (Olympus, Japan).

### Sample preparation for *in vitro* ubiquitin-dependent proteasome degradation

The human ubiquitin was prepared, as previously described^61,62^. K48-linked diubiquitin and tetra-ubiquitin were prepared following an established protocol^63^, with the conjugation reaction catalyzed with 2.5 μM human E1 and 20 μM E2-25K, in 20 mM pH 8.0 Tris-HCl buffer. Ub/K48R mutant was incorporated as the distant Ub, and Ub/77D mutant was incorporated as the proximal Ub to obtain the Ub chain of the desired length. Eventually, the additional residue D77 at the C-terminus was removed with hydrolase YUH1. The fusion protein His-TEV-Ub-GFP was prepared recombinantly based on the design previously described^40,64^. The *Pediculus humanus corporis* PINK1 kinase (phPINK1) was prepared and purified as previously described^47,65^. Ub phosphorylation was confirmed with electrospray mass spectrometry (Agilent G6530 Q-TOF).

The 26S proteasome from HEK293 cells was prepared following an established protocol, with the purification tag appended at the C-terminus of Rpn11^66,67^. The proteasome activity was assessed with a fluorogenic peptide, N-succinyl-Leu-Leu-Val-Tyr-7-amido-4-methyl coumarin (Sigma, Massachusetts, USA; S6510). A final concentration of 100 nM proteasome and 100 μM proteasome peptide substrate was prepared in the reaction buffer, containing 50 mM Tris-HCl pH 7.5, 4 mM ATP (Sigma-Aldrich, Cat# A6559), and 5 mM MgCl_2_. The fluorescence intensity of the product was measured by a fluorometer (Horiba Scientific, FluoroMax-4) at an excitation wavelength of 360 nm.

Western blotting was used to assess the degradation rate of K48-polyUb-GFP and pK48-polyUb-GFP. 200 nM proteasome substrate and 10 nM human 26S proteasome were prepared in 20 mM Tris pH 8.0 buffer, with 50 mM NaCl, 5 mM ATP, and 5 mM MgCl_2_. The mixture was incubated at 37 ° C for 0, 10, 30, 60, 90, and 120 min. At each time point, 20 µL samples were taken for Western blotting analysis on the NC membrane (Millipore, 0.45 µm). A mouse anti-GFP antibody (1:2000, San-Ying Proteintech Group, 66002-1) and an HRP-conjugated goat anti-mouse IgG (1:3000, CST, Pennsylvania, USA; 7076S) were used, with 10 s exposure. ImageJ was used to analyze the Western blot band intensities. All bands were normalized to the intensity of the GFP band. Three biological repeats were performed. Two-way ANOVA was performed using Prism software, with the phosphorylated protein band intensity (the percentage of GFP protein remaining) compared to the band intensity at time 0, or the band intensity of the unphosphorylated protein at the same time point, using two-way ANOVA in the Prism software, followed with Newman-Keuls multiple comparisons test.

### *In vitro* ubiquitin chain formation

The recombinant Ub, pUb (phosphorylated by phPINK1), Ub/S65A, and Ub/S65E were reacted with 2.5 μM human E1 and 20 μM E2-25K in 20 mM pH 8.0 Tris-HCl buffer. The formed Ub chain was determined by Coomassie blue staining on SDS page gel.

### TIRF analysis of ubiquitin-proteasome interaction

The coverslips were prepared following an established protocol^68^. Each coverslip was divided into multiple lanes with double-sided tape for parallel TIRF experiments, thus ensuring the repeatability with different proteins added to the same slide. Streptavidin (VWR Life Science, Cat# 97062-808) in imaging buffer at a concentration of 50 µg/ml was loaded; excess unbound streptavidin was washed away. The imaging buffer contains 25 mM Tris-HCl (pH 7.5), 50 mM NaCl, 10 mM MgCl2, 40 mM imidazole, 5 m/mL BSA (Sigma-Aldrich), 2.5 mM ADP (Sigma-Aldrich), and 0.5 mM ATP-γ-S (Sigma-Aldrich), as previously described^42^.

The purified human 26S proteasome at 20 nM in the imaging buffer was added to the streptavidin-immobilized coverslip, with excess unbound proteins washed away with the imaging buffer. The substrate protein, either K48-polyUb-GFP or pK48-polyUb-GFP, was then added to the coverslip to a final concentration of 400 pM. The imaging was performed on a Nikon A1 TIRF microscope equipped with a 488 nm laser and an EMCCD camera (Andor DU-897). The time series were acquired at 300 ms per frame for a total duration of 2 min for each time series. The view area was 512 pixels by 512 pixels (16 by 16 µm), and the experiments were repeated four times on four different proteasome-immobilized cover slides. The puncta density was counted in iSMS software^69^ with default settings. In particular, (1) only when the substrate protein becomes bound to the immobilized proteasome, the green puncta can be observed; (2) only the puncta with brightness above a certain threshold (≥ 3000 a.u.) and the dwell time of more than three consecutive frames are selected for counting the overall number of puncta. The dwell times were tabulated in 300 ms bins and fitted with single exponential decay functions.

### Peptide enrichment and analysis by LC-MS

Hippocampus samples were rinsed using PBS and lysed in DOC lysis buffer (1% SDS, 100 mM Tris pH 8.8, 10 mM TCEP, 40 mM CAA). Then, the samples were denatured at 95 °C for 5 min, followed by sonication at 4 °C (3 s on, 3 s off, at amplitude 30%). The lysate was centrifuged at 16000g for 10 min at 4 °C. The supernatant was collected as whole tissue extract. Protein concentration was determined by Bradford protein assay.

Extracts from the protein sample (2 g) were digested with trypsin. Then, 15 mg TiO_2_-coupled beads were incubated with 500 μl binding buffer (BB) at room temperature for 10 min. Separate TiO_2_ into three 1.5 ml EP equally, 5 mg each, and centrifuge 2000 G for 2min. Resolve digested peptides with 600 μl BB and combine with 5 mg incubated TiO_2_ at room temperature for 30 min and centrifuge 1000 G for 2 min. Repeat the solved procedures for the dried peptides twice with 5 mg incubated TiO_2_ and discard the supernatant. Peptides were eluted using acetonitrile 40% at pH 10. Then the elution was dried in a vacuum concentrator.

The enriched peptides sample were resolved in solvent A (0.1% formic acid). After centrifuging at 16000 G for 10 min, the peptides in the supernatant were sequenced using Orbitrap Fusion mass spectrometers (Thermo Fisher Scientific, Rockford, IL, US) coupled with an Easy-nLC 1000 nanoflow LC system (Thermo Fisher Scientific). The injected peptides were separated on a reverse-phase nano-HPLC C18 column (Precolumn, 3 μm, 120 Å, 2X100 i.d.; analyzed column, 1.9 μm 120 Å, 30 cm X 150 μm, i.d.) at a flow rate of 600 nL/min with a 150 min gradient of 7-95% solvent B (0.1% formic acid in acetonitrile).

For peptide ionization, 2100 V was applied, and a 320 °C capillary temperature was used. For detection with Q-Exactive HF, a precursor scan was carried out in the Orbitrap by scanning m/z 350-1400 with a resolution of 120,000 at 200 m/z. The most intense ion selected under top-speed mode were isolated in Quadrupole with a 1.6 m/z window and fragmented by HCD with a normalized collision energy of 27%, then measured in the linear ion trap using the rapid ion trap scan rate. Automatic gain control targets were 5 ×10^4^ ions with a max injection time of 19 ms for full scans and 5×10^3^ ions with 35 ms for MS/MS scans. Dynamic exclusion time was set as 30 sec.

The MS analysis for QE HF was performed with one full scan (350–1400LJm/z, RLJ=LJ120,000 at 200LJm/z) at an automatic gain control target of 3×10^6^ ions, max injection time 80 ms, followed by up to 30 data-dependent MS/MS scans with higher-energy collision dissociation (target 5×10^4^LJions, max injection time 19 ms, isolation window 1.6LJm/z, normalized collision energy of 27%), detected in the Orbitrap (RLJ=LJ15,000 at 200LJm/z). Dynamic exclusion time was set as 30 s.

### MS data processing

Raw files were searched against the mouse National Center for Biotechnology Information (NCBI) Refseq protein database (updated on 2013/07/01, 29764 entries) by the software Thermo Proteome Discoverer 2.1 using Mascot 2.3. The mass tolerances were 20 ppm for precursor and 50 m.mu for product ions for Fusion. The search engine set acetyl (protein N-term), oxidation(M), Phospho(S/T), Phospho(Y) as variable modification: acetyl (protein N-term), Oxidation(M), Phospho(ST), Phospho(Y). Trypsin total digestion of up to two missed cleavages was allowed. The peptide identifications were accepted at a false discovery rate (FDR) of 1%. Label-free protein quantifications were calculated using a label-free approach. A unique peptide was used to represent the absolute abundance of a particular peptide across the sample.

### Proteomics data analysis

Quantization normalization was applied to remove the batch effect. Missing values were filled with the minimum value for the benefit of the following analysis. The differentially expressed proteins were defined as meeting the following two criteria: 1) Fold change value less than 0.5 or more than 2, 2) with top 75 expression accounts in the proteome. Gene Set Enrichment Analysis (GSEA) - based Gene Ontology (GO) analysis was conducted by GSEApy (version 0.10.5) Python package to find enriched GO terms. The e of proteins to GO terms were derived from DAVID website (2021 update). The ranking metric of GSEA was the fold change, and the cutoff P-value was 0.05 to filter the significantly enriched GO terms.

### Statistical analysis

Data are presented as mean ± SD or media with 95% confidence intervals (CI). GraphPad Prism Software (version 6.0, GraphPad Software Inc., San Diego, CA, USA) was used for statistical analysis and plot graphs. The ROUT test was used to identify outliers. The Brown-Forsythe test was used to evaluate the equal variances of the data. D’Agonstino-Pearson omnibus normality test was performed to test the normality of the data. If the data pass the normality test and equal variance test, we used the *t*-test or the parametric one-way ANOVA (Tukey multiple comparisons test) or two-way ANOVA (Newman-Keuls multiple comparisons test). If the data does not pass the normality test, we use the nonparametric test (Dunn’s multiple comparisons test). We use the Chi-square test to assess the difference between the categorical data. A value of *P* < 0.05 was considered statistically significant.

## Acknowledgements

This study was supported by the National Key R&D Program of China (2018YFA0507700) to C.T. and (2021YFF1200900) to T.T.L., the National Natural Science Foundation of China (81971304) to W.P.Z. and (32070666) to T.T.L.. The funders had no role in study design, data collection and analysis, decision to publish or preparation of the manuscript.

## Author contributions

W.P.Z. and C.T. designed the study and wrote the manuscript. C.C., H.W.Y., Y.Z., T.W., T.Y.G., Z.L.L. and T.F.W. performed the experiments. Y.B.L. and W.P.Z. analyzed the data at mouse and cellular level. T.T.L. analyzed the proteomic data. C.T. analyzed the data at protein level.

## Competing interests

The authors declare that they have no competing interests.

**Supplementary information** includes 7 figures.

**Correspondence and requests for material** should be addressed to Wei-Ping Zhang or Chun Tang

## Notes

### Competing Interest Statement

The authors have declared no competing interest.

## References

1. Lopez-Otin, C., Blasco, M.A., Partridge, L., Serrano, M. & Kroemer, G. Hallmarks of aging: An expanding universe. Cell 186, 243–278 (2023).

2. Wang, Q., et al. Role of mitophagy in the neurodegenerative diseases and its pharmacological advances: A review. Front Mol Neurosci 15, 1014251 (2022).

3. Bulteau, A.L., Petropoulos, I. & Friguet, B. Age-related alterations of proteasome structure and function in aging epidermis. Exp. Gerontol. 35, 767–777 (2000).

4. Keller, J.N., Hanni, K.B. & Markesbery, W.R. Possible involvement of proteasome inhibition in aging: implications for oxidative stress. Mech Ageing Dev 113, 61–70 (2000).

5. Petropoulos, I., et al. Increase of oxidatively modified protein is associated with a decrease of proteasome activity and content in aging epidermal cells. J. Gerontol. A Biol. Sci. Med. Sci. 55, B220–227 (2000).

6. Davidson, K. & Pickering, A.M. The proteasome: A key modulator of nervous system function, brain aging, and neurodegenerative disease. Front Cell Dev Biol 11, 1124907 (2023).

7. Schmidt, M.F., Gan, Z.Y., Komander, D. & Dewson, G. Ubiquitin signalling in neurodegeneration: mechanisms and therapeutic opportunities. Cell Death Differ. 28, 570–590 (2021).

8. Hegde, A.N., Goldberg, A.L. & Schwartz, J.H. Regulatory subunits of cAMP-dependent protein kinases are degraded after conjugation to ubiquitin: a molecular mechanism underlying long-term synaptic plasticity. Proc Natl Acad Sci U S A 90, 7436–7440 (1993).

9. Palombella, V.J., Rando, O.J., Goldberg, A.L. & Maniatis, T. The ubiquitin-proteasome pathway is required for processing the NF-kappa B1 precursor protein and the activation of NF-kappa B. Cell 78, 773–785 (1994).

10. Gouet, C., Aburto, B., Vergara, C. & Sanhueza, M. On the mechanism of synaptic depression induced by CaMKIIN, an endogenous inhibitor of CaMKII. PloS one 7, e49293 (2012).

11. Munkacsy, E., et al. Neuronal-specific proteasome augmentation via Prosbeta5 overexpression extends lifespan and reduces age-related cognitive decline. Aging Cell 18, e13005 (2019).

12. Lowe, J., et al. Ubiquitin is a common factor in intermediate filament inclusion bodies of diverse type in man, including those of Parkinson’s disease, Pick’s disease, and Alzheimer’s disease, as well as Rosenthal fibres in cerebellar astrocytomas, cytoplasmic bodies in muscle, and mallory bodies in alcoholic liver disease. J. Pathol. 155, 9–15 (1988).

13. Alves-Rodrigues, A., Gregori, L. & Figueiredo-Pereira, M.E. Ubiquitin, cellular inclusions and their role in neurodegeneration. Trends Neurosci. 21, 516–520 (1998).

14. Komander, D. & Rape, M. The ubiquitin code. Annu. Rev. Biochem. 81, 203–229 (2012).

15. Shi, Y., et al. Rpn1 provides adjacent receptor sites for substrate binding and deubiquitination by the proteasome. Science 351(2016).

16. Deveraux, Q., Ustrell, V., Pickart, C. & Rechsteiner, M. A 26 S protease subunit that binds ubiquitin conjugates. J. Biol. Chem. 269, 7059–7061 (1994).

17. Husnjak, K., et al. Proteasome subunit Rpn13 is a novel ubiquitin receptor. Nature 453, 481–488 (2008).

18. Swatek, K.N. & Komander, D. Ubiquitin modifications. Cell Res 26, 399–422 (2016).

19. Valente, E.M., et al. Hereditary early-onset Parkinson’s disease caused by mutations in PINK1. Science 304, 1158–1160 (2004).

20. Koyano, F., et al. Ubiquitin is phosphorylated by PINK1 to activate parkin. Nature 510, 162–166 (2014).

21. Lazarou, M., et al. The ubiquitin kinase PINK1 recruits autophagy receptors to induce mitophagy. Nature 524, 309–314 (2015).

22. Okatsu, K., Kimura, M., Oka, T., Tanaka, K. & Matsuda, N. Unconventional PINK1 localization to the outer membrane of depolarized mitochondria drives Parkin recruitment. J. Cell Sci. 128, 964–978 (2015).

23. Valente, E.M., et al. Localization of a novel locus for autosomal recessive early-onset parkinsonism, PARK6, on human chromosome 1p35-p36. Am. J. Hum. Genet. 68, 895–900 (2001).

24. Jin, S.M., et al. Mitochondrial membrane potential regulates PINK1 import and proteolytic destabilization by PARL. J. Cell Biol. 191, 933–942 (2010).

25. Greene, A.W., et al. Mitochondrial processing peptidase regulates PINK1 processing, import and Parkin recruitment. EMBO reports 13, 378–385 (2012).

26. Yamano, K. & Youle, R.J. PINK1 is degraded through the N-end rule pathway. Autophagy 9, 1758–1769 (2013).

27. Gao, J., et al. Cytosolic PINK1 promotes the targeting of ubiquitinated proteins to the aggresome-autophagy pathway during proteasomal stress. Autophagy 12, 632–647 (2016).

28. Takatori, S., Ito, G. & Iwatsubo, T. Cytoplasmic localization and proteasomal degradation of N-terminally cleaved form of PINK1. Neurosci Lett 430, 13–17 (2008).

29. Wauer, T., et al. Ubiquitin Ser65 phosphorylation affects ubiquitin structure, chain assembly and hydrolysis. EMBO J 34, 307–325 (2015).

30. Fiesel, F.C., et al. (Patho-)physiological relevance of PINK1-dependent ubiquitin phosphorylation. EMBO reports 16, 1114–1130 (2015).

31. Hou, X., et al. Age- and disease-dependent increase of the mitophagy marker phospho-ubiquitin in normal aging and Lewy body disease. Autophagy 14, 1404–1418 (2018).

32. Hou, X., et al. Alpha-synuclein-associated changes in PINK1-PRKN-mediated mitophagy are disease context dependent. Brain Pathol., e13175 (2023).

33. Hakim, V., Cohen, L.D., Zuchman, R., Ziv, T. & Ziv, N.E. The effects of proteasomal inhibition on synaptic proteostasis. EMBO J 35, 2238–2262 (2016).

34. Thibaudeau, T.A., Anderson, R.T. & Smith, D.M. A common mechanism of proteasome impairment by neurodegenerative disease-associated oligomers. Nat Commun 9, 1097 (2018).

35. Wang, L., et al. PTEN-L is a novel protein phosphatase for ubiquitin dephosphorylation to inhibit PINK1-Parkin-mediated mitophagy. Cell Res 28, 787–802 (2018).

36. Wall, C.E., et al. PPEF2 Opposes PINK1-Mediated Mitochondrial Quality Control by Dephosphorylating Ubiquitin. Cell Rep 29, 3280–3292 e3287 (2019).

37. Ye, S.-X., et al. PP2A forestalls mitophagy by dephosphorylating Parkin and ubiquitin. bioRxiv (2022). doi: https://doi.org/10.1101/2022.03.12.484070

38. Ordureau, A., et al. Quantitative proteomics reveal a feedforward mechanism for mitochondrial PARKIN translocation and ubiquitin chain synthesis. Mol Cell 56, 360–375 (2014).

39. Beilina, A., et al. Mutations in PTEN-induced putative kinase 1 associated with recessive parkinsonism have differential effects on protein stability. Proc Natl Acad Sci U S A 102, 5703–5708 (2005).

40. Dantuma, N.P., Lindsten, K., Glas, R., Jellne, M. & Masucci, M.G. Short-lived green fluorescent proteins for quantifying ubiquitin/proteasome-dependent proteolysis in living cells. Nat Biotechnol 18, 538–543 (2000).

41. Samant, R.S., Livingston, C.M., Sontag, E.M. & Frydman, J. Distinct proteostasis circuits cooperate in nuclear and cytoplasmic protein quality control. Nature 563, 407–411 (2018).

42. Lu, Y., Lee, B.H., King, R.W., Finley, D. & Kirschner, M.W. Substrate degradation by the proteasome: a single-molecule kinetic analysis. Science 348, 1250834 (2015).

43. Li, J., et al. Proteome-wide mapping of short-lived proteins in human cells. Mol Cell 81, 4722–4735 e4725 (2021).

44. Alkalay, I., et al. Stimulation-dependent I kappa B alpha phosphorylation marks the NF-kappa B inhibitor for degradation via the ubiquitin-proteasome pathway. Proc Natl Acad Sci U S A 92, 10599–10603 (1995).

45. Press, M., Jung, T., Konig, J., Grune, T. & Hohn, A. Protein aggregates and proteostasis in aging: Amylin and beta-cell function. Mech Ageing Dev 177, 46-54 (2019).

46. Pollock, L., Jardine, J., Urbe, S. & Clague, M.J. The PINK1 repertoire: Not just a one trick pony. Bioessays 43, e2100168 (2021).

47. Wauer, T., Simicek, M., Schubert, A. & Komander, D. Mechanism of phospho-ubiquitin-induced PARKIN activation. Nature 524, 370–374 (2015).

48. Bard, J.A.M., Bashore, C., Dong, K.C. & Martin, A. The 26S Proteasome Utilizes a Kinetic Gateway to Prioritize Substrate Degradation. Cell 177, 286-+ (2019).

49. Trigo, D., Nadais, A. & da Cruz, E.S.O.A.B. Unravelling protein aggregation as an ageing related process or a neuropathological response. Ageing Res Rev 51, 67–77 (2019).

50. Suresh, K., Mattern, M., Goldberg, M.S. & Butt, T.R. The Ubiquitin Proteasome System as a Therapeutic Area in Parkinson’s Disease. Neuromolecular Med. (2023).

51. Devi, L., Prabhu, B.M., Galati, D.F., Avadhani, N.G. & Anandatheerthavarada, H.K. Accumulation of amyloid precursor protein in the mitochondrial import channels of human Alzheimer’s disease brain is associated with mitochondrial dysfunction. J Neurosci 26, 9057–9068 (2006).

52. Kopeikina, K.J., et al. Tau accumulation causes mitochondrial distribution deficits in neurons in a mouse model of tauopathy and in human Alzheimer’s disease brain. Am J Pathol 179, 2071–2082 (2011).

53. Imaizumi, Y., et al. Mitochondrial dysfunction associated with increased oxidative stress and alpha-synuclein accumulation in PARK2 iPSC-derived neurons and postmortem brain tissue. Mol Brain 5, 35 (2012).

54. Hamilton, A.M., et al. Activity-dependent growth of new dendritic spines is regulated by the proteasome. Neuron 74, 1023–1030 (2012).

55. Rodriguez-Ortiz, C.J., Balderas, I., Saucedo-Alquicira, F., Cruz-Castaneda, P. & Bermudez-Rattoni, F. Long-term aversive taste memory requires insular and amygdala protein degradation. Neurobiol. Learn. Mem. 95, 311–315 (2011).

56. Veeranna, et al. Declining phosphatases underlie aging-related hyperphosphorylation of neurofilaments. Neurobiol. Aging 32, 2016–2029 (2011).

57. Xing, J., et al. Protein phosphatase 2A activators reverse age-related behavioral changes by targeting neural cell senescence. Aging Cell 22, e13780 (2023).

58. Cong, L., et al. Multiplex genome engineering using CRISPR/Cas systems. Science 339, 819–823 (2013).

59. Shen, C., et al. The Depletion of NAMPT Disturbs Mitochondrial Homeostasis and Causes Neuronal Degeneration in Mouse Hippocampus. Molecular neurobiology 60, 1267–1280 (2023).

60. Ran, F.A., et al. Genome engineering using the CRISPR-Cas9 system. Nat. Protoc. 8, 2281–2308 (2013).

61. Liu, Z., et al. Lys63-linked ubiquitin chain adopts multiple conformational states for specific target recognition. eLife 4, e05767 (2015).

62. Liu, Z., et al. Noncovalent dimerization of ubiquitin. Angewandte Chemie. International Ed. In English 51, 469-472 (2012).

63. Pickart, C.M. & Raasi, S. Controlled synthesis of polyubiquitin chains. Methods Enzymol 399, 21–36 (2005).

64. Inobe, T., Fishbain, S., Prakash, S. & Matouschek, A. Defining the geometry of the two-component proteasome degron. Nature chemical biology 7, 161–167 (2011).

65. Dong, X., et al. Ubiquitin S65 phosphorylation engenders a pH-sensitive conformational switch. Proc Natl Acad Sci U S A 114, 6770–6775 (2017).

66. Huang, X., Luan, B., Wu, J. & Shi, Y. An atomic structure of the human 26S proteasome. Nat Struct Mol Biol 23, 778–785 (2016).

67. Guerrero, C., Tagwerker, C., Kaiser, P. & Huang, L. An integrated mass spectrometry-based proteomic approach: quantitative analysis of tandem affinity-purified in vivo cross-linked protein complexes (QTAX) to decipher the 26 S proteasome-interacting network. Mol Cell Proteomics 5, 366–378 (2006).

68. Roy, R., Hohng, S. & Ha, T. A practical guide to single-molecule FRET. Nat Methods 5, 507–516 (2008).

69. Preus, S., Noer, S.L., Hildebrandt, L.L., Gudnason, D. & Birkedal, V. iSMS: single-molecule FRET microscopy software. Nat Methods 12, 593–594 (2015).

