## Supplemental Figures for "Ubiquitin phosphorylation accelerates protein aggregation and promotes neurodegeneration in the aging brain"

**Short title:** Elevated phosphorylated ubiquitin linked to neurodegeneration

**Supplementary figures**

**Includes 7 figures**

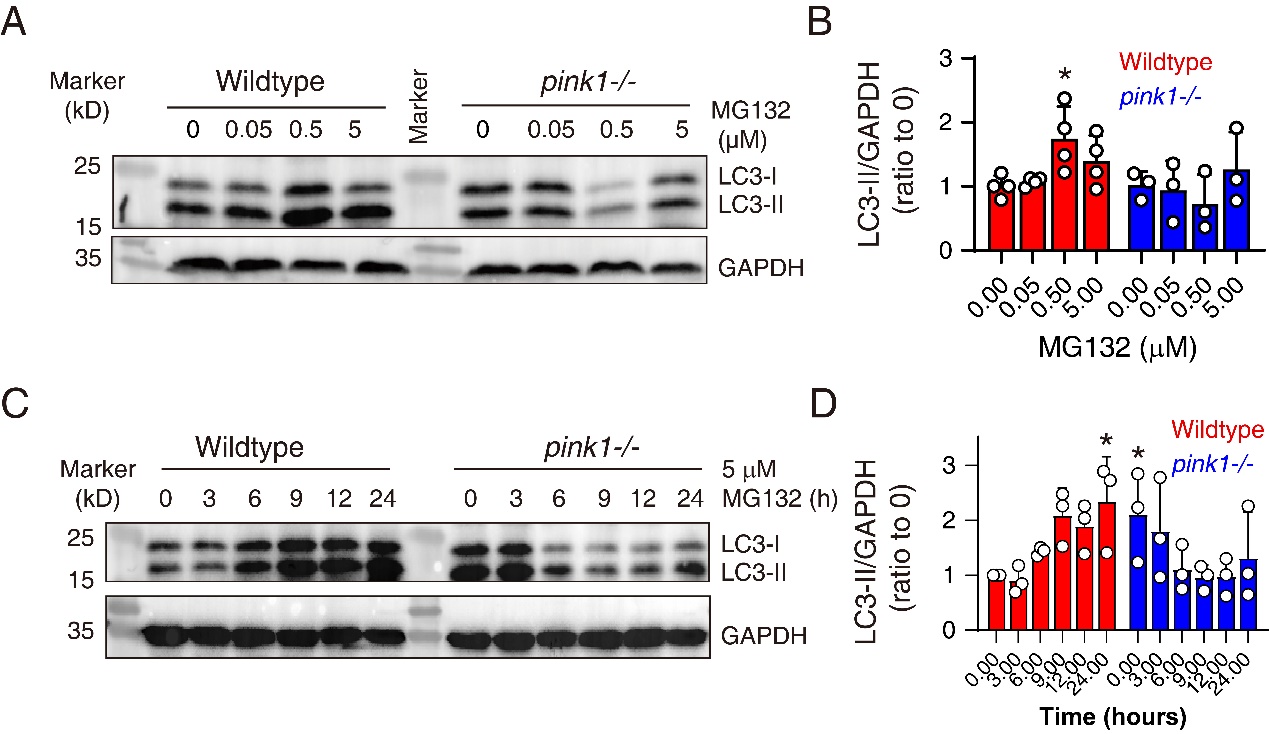

**Figure S1** **The knockout of *pink1* gene inhibited MG132-induced autophagy in HEK293 cells. A, B**. Western blot analysis of LC3 at 8 hours after the administration of MG132 in wildtype and *pink1-/-* cells. *N*=4. **P*<0.05, compared with “0” MG132, one-way ANOVA. **C, D**. Western blot analysis of LC3 at 0-24 hours after the administration of MG132 in wildtype and *pink1-/-* cells. *N*=4. **P*<0.05, compared with time 0, one-way ANOVA.

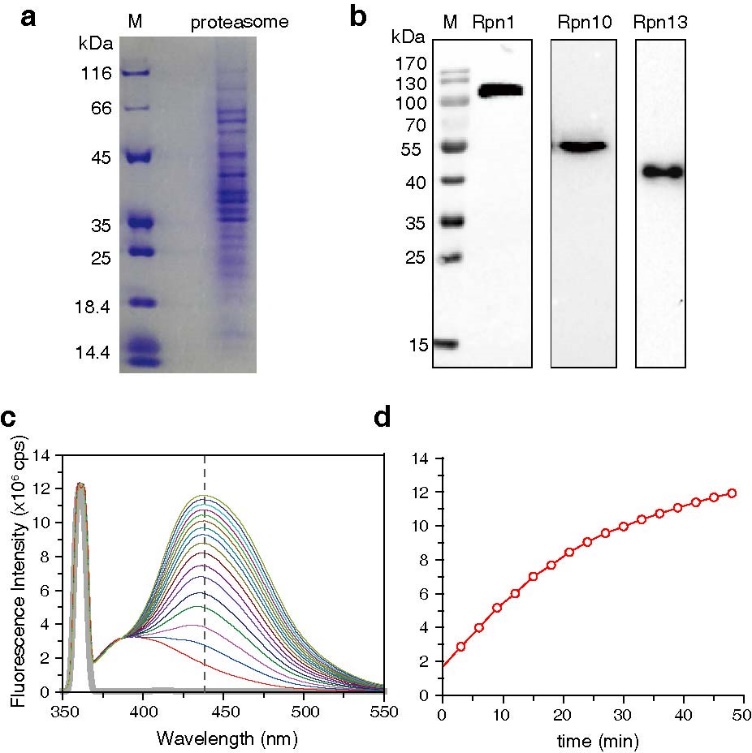

**Fig. S2 Assessment of 26S proteasome activity with protein *in vitro* degradation. A**. The Coomassie blue staining of proteins composing proteasome. **B.** Western blot identification of the three Ub receptors in the 26S proteasome, including Rpn1, Rpn10, and Rpn13. **C.** The determination of *in vitro* proteasome degradation activity using a fluorogenic peptide.

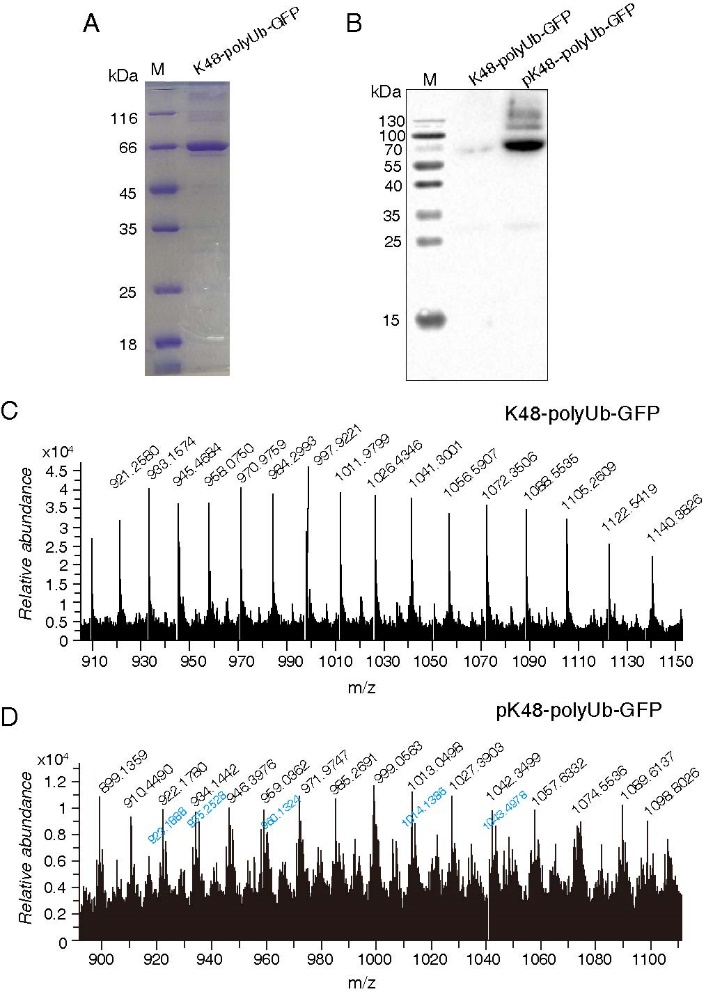

**Fig. S3** **Identification of the K48-polyUb-GFP and pK48-polyUb-GFP as the proteasomal substrates.** K48-linked tetraubiquitin is covalent linked to Ub-GFP fusion protein, also via a K48 linkage. **A**. The identity of the substrate protein was assessed with SDS-PAGE. **B**. Phosphorylation of the substrate protein by sPINK1 was confirmed with Western blot using an anti-pUb antibody. **C, D**. ESI-MS analysis of K48-polyUb-GFP (C) and pK48-polyUb-GFP (D). The m/z profile indicates that the protein carries 1-2 phosphoryl groups, corresponding to black and blue labels, respectively.

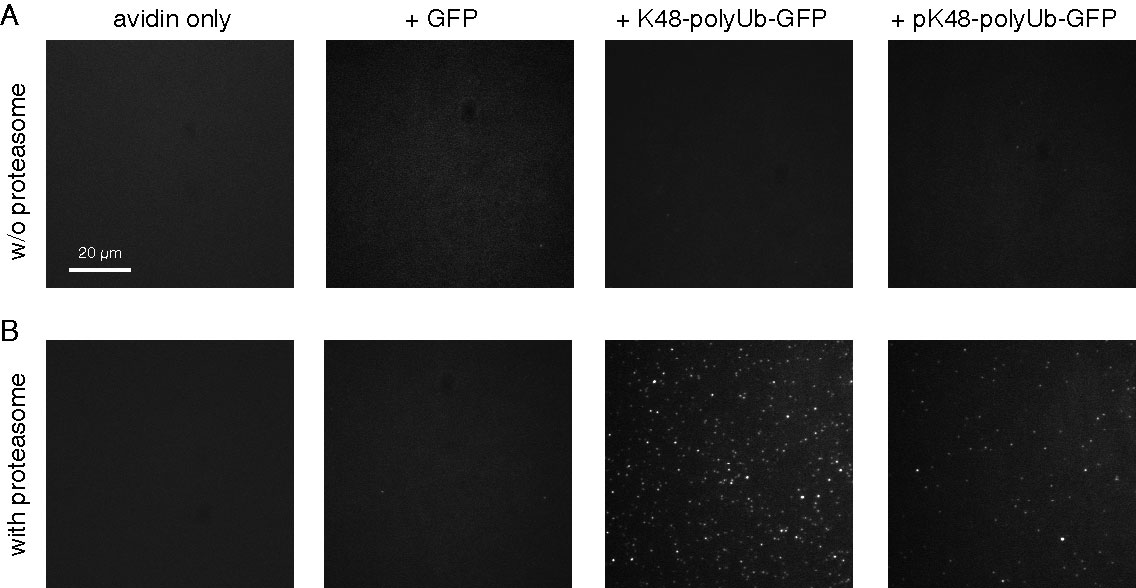

**Fig. S4 Visualization of proteasome-associated substrate protein using TIRF microscopy.** **A**. The representative images of TIRF without the 26S proteasome immobilized at the coverslip. **B**. The representative images of TIRF with the 26S proteasome immobilized at the coverslip. The immobilized proteasome alone or the application of GFP protein alone cannot be visualized using TIRF. The K48-polyUb-GFP can be visualized with TIRF as discrete puncta upon binding to the immobilized proteasome.

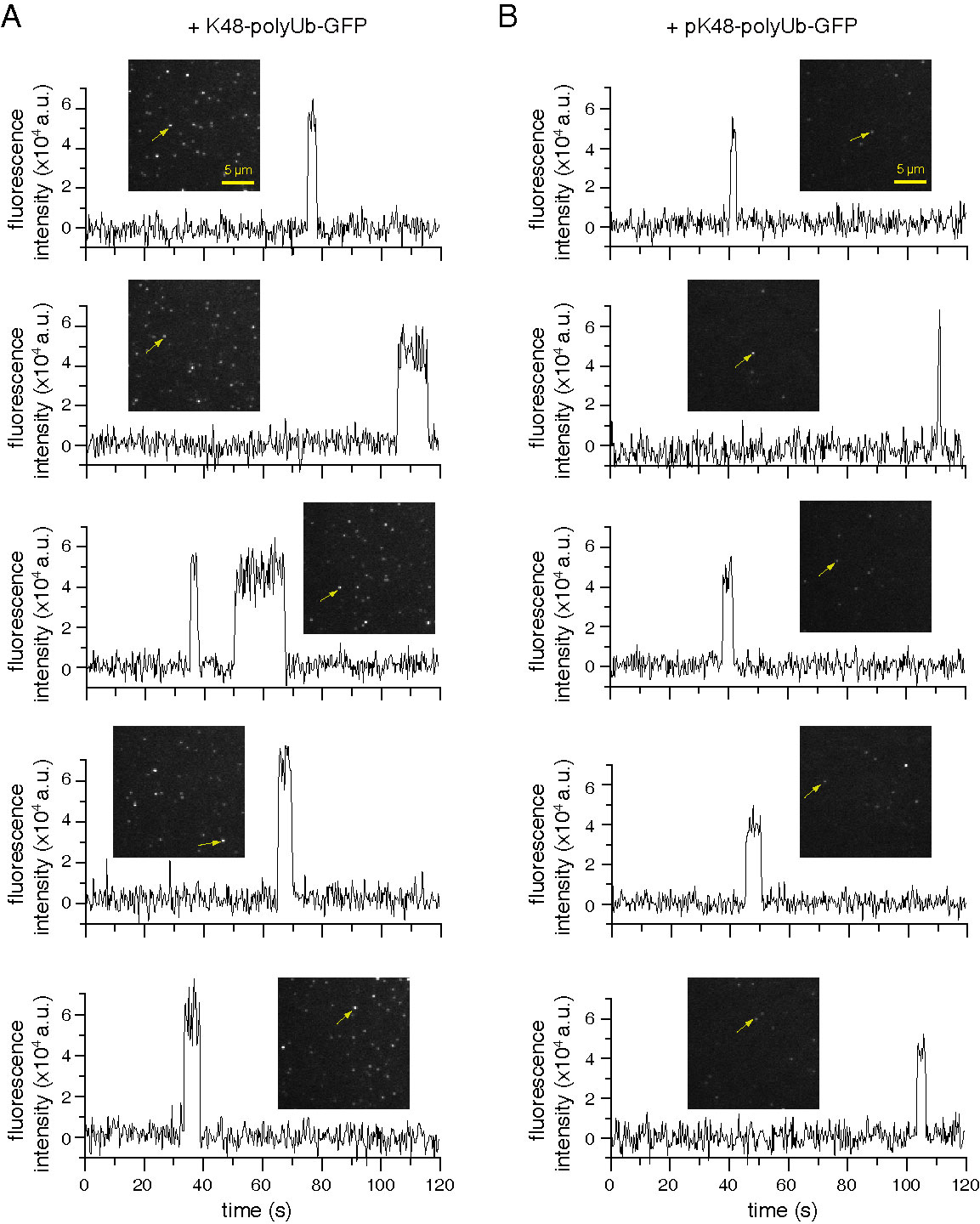

**Fig. S5 Ub phosphorylation accelerated the dissociation of K48-polyUb-GFP from the immobilized proteasome.** From top to bottom, five representative time traces for K48-polyUb-GFP (A) and pK48-polyUb-GFP (B), respectively, associated with the immobilized 26S proteasome. The analyzed puncta are indicated with an arrow.

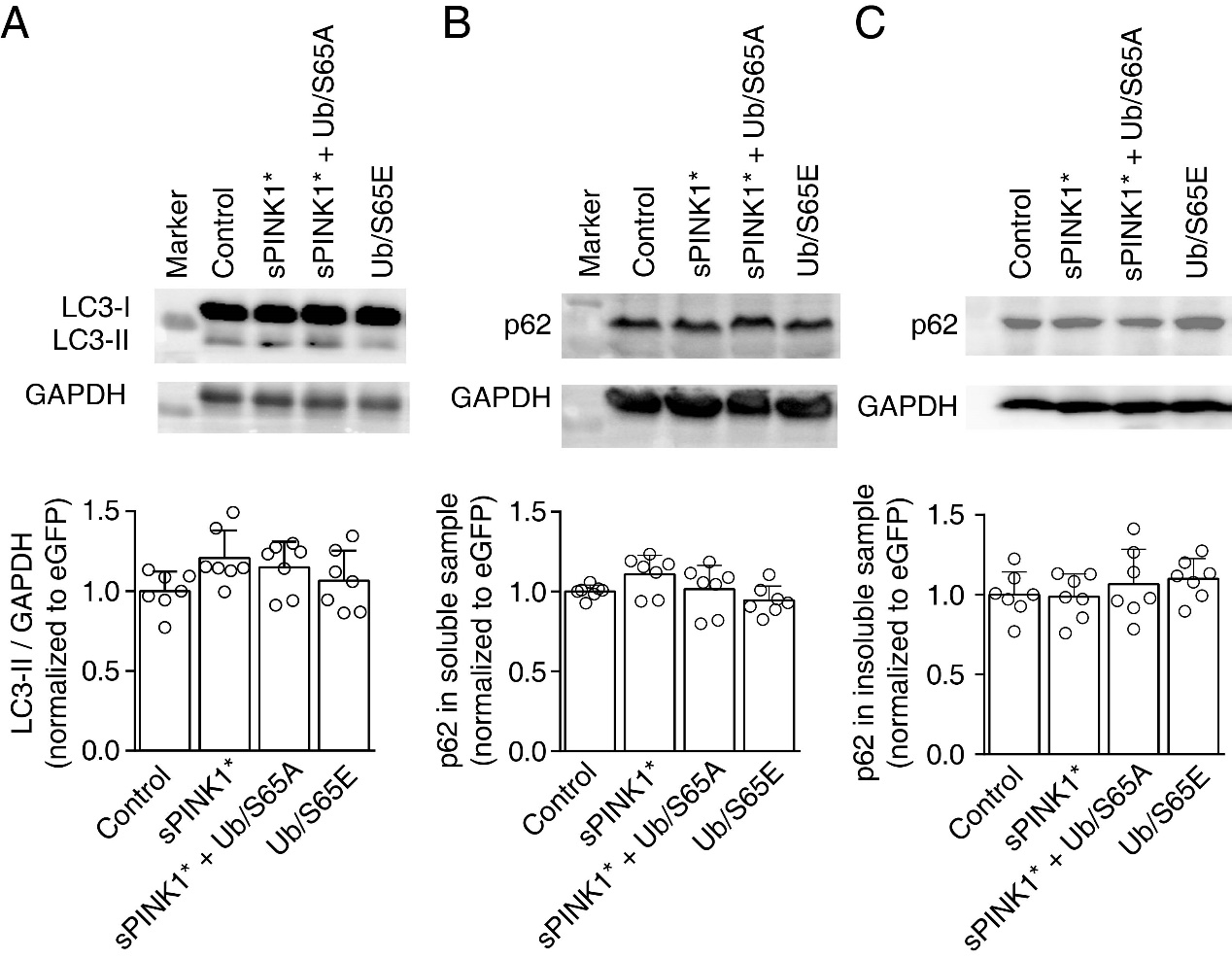

**Fig. S6 The elevation of pUb level in the mouse hippocampus neurons did not change the autophagy flow.** Mice were sacrificed on 70^th^ day post AAV2/9 injection. The protein sample was separated as soluble sample (A and B) and insoluble sample (C). **A.** The level of LC3 in the soluble sample. **B, C**. The level of p62 in the soluble sample (B) and the insoluble sample (C). The upper panel are the representative graphs of the Western blot, and the lower level are the corresponding statistical analysis.

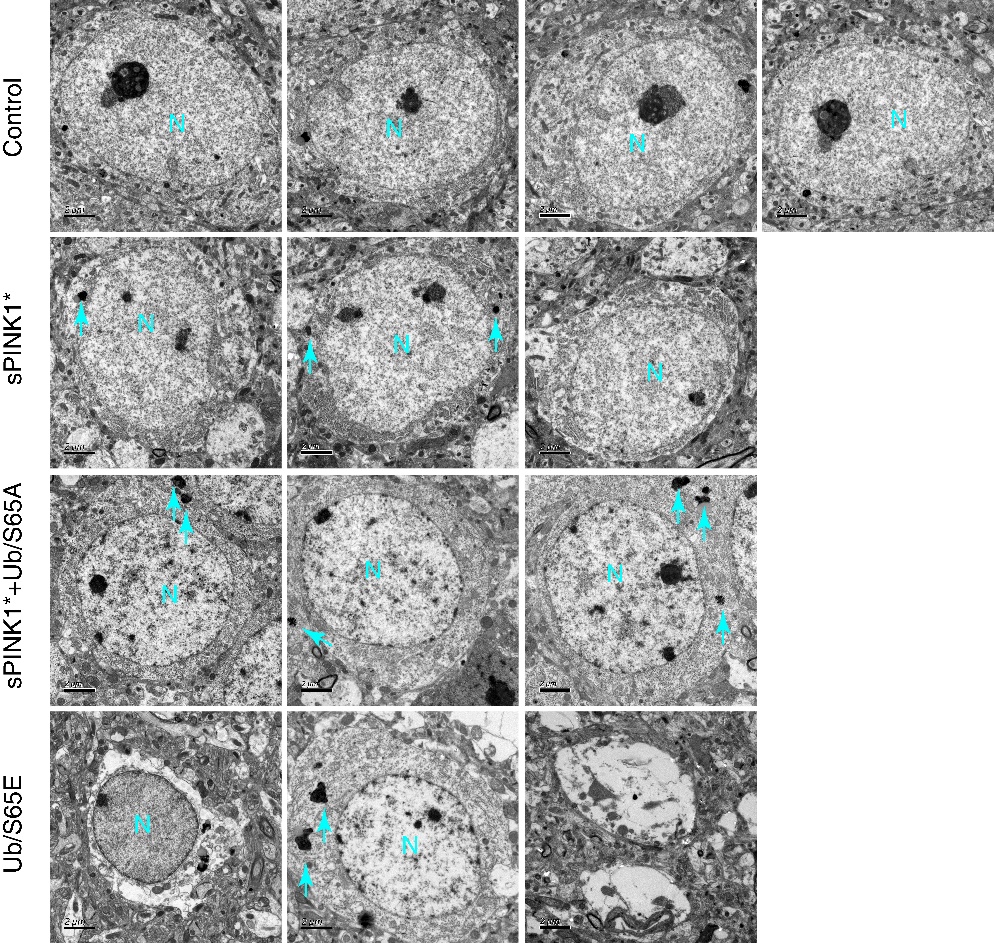

**Fig. S7 Representative graphs of transmission electron microscope of the hippocampus neurons.** Mice were sacrificed on 70^th^ day post AAV2/9 injection. N denotes the nucleus.
